# Topology and habitat assortativity drive neutral and adaptive diversification in spatial graphs

**DOI:** 10.1101/2021.07.06.451404

**Authors:** Victor Boussange, Loïc Pellissier

## Abstract

Biodiversity results from differentiation mechanisms developing within biological populations. Such mechanisms are influenced by the properties of the landscape over which individuals interact, disperse and evolve. Notably, landscape connectivity and habitat heterogeneity constrain the movement and survival of individuals, thereby promoting differentiation through drift and local adaptation. Nevertheless, the complexity of landscape features can blur our understanding of how they drive differentiation. Here, we formulate a stochastic, eco-evolutionary model where individuals are structured over a graph that captures complex connectivity patterns and accounts for habitat heterogeneity. Individuals possess neutral and adaptive traits, whose divergence results in differentiation at the population level. The modelling framework enables an analytical underpinning of emerging macroscopic properties, which we complement with numerical simulations to investigate how the graph topology and the spatial habitat distribution affect differentiation. We show that in the absence of selection, graphs with high characteristic length and high heterogeneity in degree promote neutral differentiation. Habitat assortativity, a metric that captures habitat spatial autocorrelation in graphs, additionally drives differentiation patterns under habitat-dependent selection. While assortativity systematically amplifies adaptive differentiation, it can foster or depress neutral differentiation depending on the migration regime. By formalising the eco-evolutionary and spatial dynamics of biological populations in complex landscapes, our study establishes the link between landscape features and the emergence of diversification, contributing to a fundamental understanding of the origin of biodiversity gradients.

**Significance statement:** It is not clear how landscape connectivity and habitat heterogeneity influence differentiation in biological populations. To obtain a mechanistic understanding of underlying processes, we construct an individual-based model that accounts for eco-evolutionary and spatial dynamics over graphs. Individuals possess both neutral and adaptive traits, whose co-evolution results in differentiation at the population level. In agreement with empirical studies, we show that characteristic length, heterogeneity in degree and habitat assortativity drive differentiation. By using analytical tools that permit a macroscopic description of the dynamics, we further link differentiation patterns to the mechanisms that generate them. Our study provides support for a mechanistic understanding of how landscape features affect diversification.

## 1 Introduction

Biodiversity results from differentiation processes influenced by the features of the landscape over which populations are distributed [1, 2]. The documentation of high levels of species diversity in mountain region or riverine systems suggests that complex connectivity patterns and habitat heterogeneity foster diversity [3, 4, 5, 6, 7]. Correlative studies support this hypothesis, establishing concertedly that connectivity and habitat heterogeneity systematically emerge as core predictors of species richness across a wide range of regions and taxonomic groups [8, 9, 10, 11, 12, 13]. However, hypotheses formulated based on empirical evidence should be complemented by mechanistic models to crystallise a causal understanding between processes and patterns [14]. While the number of simulation studies is growing steadily [15], such studies lack a mathematical formalism to facilitate the interpretation of the model outcomes by providing an analytical underpinning to the simulation results [16].

Differentiation processes emerge as a result of mutation, selection and migration and can be classified as neutral or adaptive [17, 18, 19]. Neutral differentiation is initiated by the stochastic drift of local phenotypes when spatial isolation and limited dispersal create barriers to gene flow, allowing distinct phenotypes to emerge in spatially structured populations [18, 20, 21]. In contrast, adaptive differentiation results from selection pressure [22, 23], which promotes distinct, locally adapted phenotypes in populations occupying patches with different habitat conditions [24, 25]. The evolution of neutral phenotypes and of adaptive phenotypes are not independent, as selective forces can indirectly select for those neutral phenotypes that happen to be linked to the fittest adaptive phenotype, a mechanism called “genetic hitchhiking” [26]. Moreover, selection pressure can generate barriers to gene flow between populations in heterogeneous habitat landscapes [27, 28, 29], a phenomenon coined “isolation by environment”, which can amplify neutral differentiation. Overall, landscape features constrain populations’ dispersal, affecting neutral and adaptive processes and their interplay. This leads to complex feedback loops that are difficult to comprehend without a formalised mechanistic model [2, 30].

Models link patterns to processes [14], and the explicit representation of the landscape within an evolutionary model can lead to a causal understanding of how landscape features shape diversity [31]. Graphs provide a convenient mathematical representation of landscapes, where vertices represent demes inhabited by populations, and edges capture the connectivity between demes [32, 33, 34, 35]. Under ecological dynamics, metapopulation models have been used to study the role of graph topology on the persistence of population [36, 37, 38, 39, 40, 41] and community diversity [42, 43, 44]. Evolutionary mechanisms are nevertheless fundamental drivers of diversity, and should therefore be explicitly integrated within models [45]. Evolutionary game theory explores how graph topology impacts the fixation probability and the fixation time of a mutated phenotype [46, 47, 48, 49, 50, 51, 52].

However, the framework does not consider the continuous accumulation of mutations, and is therefore not suited to address the emergence of diversity. By combining a metapopulation model with a model of neutral evolution, [31, 53] investigated how graph topology affects neutral diversity. This phenomenological approach demonstrated the key role of topological properties in shaping diversity, and its predictions could be matched with empirical data from e.g. river basins [54]. Nevertheless, diversity results from the combination of neutral and adaptive processes developing at the population level [55, 56, 57]. A first principles modelling approach considering graphs but also building upon the elementary processes of ecological interactions, reproduction, mutation and migration may therefore be promising to investigate emergent patterns.

Stochastic models for structured populations, rooted in the microscopic description of individuals, offer a generic framework for modelling eco-evolutionary dynamics [58, 59]. In particular, such models can capture the interplay between population dynamics, spatial dynamics and phenotypic evolution, while providing a rigorous set-up for analytical investigation. By anchoring this modelling paradigm in a mathematical framework, the work of Champagnat et al. [58] generalises models of population genetics [60, 61, 62, 63] and quantitative genetics [64, 65, 66, 67], which have early on stimulated research into the link between spatial population structure and neutral differentiation. The framework embraces density-dependent selection, which could explain the emergence of sympatric speciation from competition processes [22], and how spatial segregation can emerge as a byproduct of these adaptive processes along environmental gradients [68]. Related models have addressed the effects of landscape dynamics and habitat heterogeneity [69, 70, 71] on diversification, with mathematical insights into the dynamics [72, 73, 74]. Because it accounts for finite population size, the baseline model of Champagnat et al. can also capture neutral differentiation dynamics and therefore the coupling between neutral and adaptive processes [75, 76]. Nonetheless, the aforementioned studies considered geographical space as implicit [69, 70] or assumed regular spatial structures (regular lattices [72, 74] or continuous space [68, 73, 71]), therefore not addressing the topological complexity of landscapes. A stochastic individual-based model using graphs as a representation of the underlying landscape could help formalise the fundamental link between differentiation processes and landscape features.

A key challenge in comprehending biodiversity patterns is to understand how individual dynamics result in differentiation at the landscape level [55, 56, 57]. Here, we investigate how complex connectivity patterns and habitat heterogeneity affect both neutral and adaptive differentiation by constructing an individual-based model (IBM) that accounts for eco-evolutionary and spatial dynamics. In this modelling approach, individuals are spatially structured over a graph and possess co-evolving neutral and adaptive traits. The finite size of local populations generates neutral differentiation by inducing a stochastic drift in the neutral trait evolution, while heterogeneous selection pressure gives rise to adaptive differentiation. Macroscopic properties of the model are analytically tractable, and we obtain a deterministic approximation of population and adaptive trait dynamics which connects the emerging patterns to the graph properties that generate them. However, neutral differentiation is stochastic by nature, which complicates its analytical underpinning. We therefore rely on numerical simulations of the IBM to measure the effect of graph topology on neutral differentiation. In the case where selection is absent (setting (1)), we investigate which graph topology metrics [77] affect neutral diversity. Second, in the case of heterogeneous selection pressure (setting (2)), we investigate the extent to which such topological properties, in combination with the spatial habitat distribution, affect levels of (i) adaptive and (ii) neutral differentiation. We expect to identify metrics that capture the effects of graph topology and habitat spatial distribution on differentiation, with the intention of relating them to the underlying processes. Overall, our study establishes the link between landscape features and population differentiation and contributes to a fundamental understanding of how landscape topology shapes biodiversity patterns.

## 2 Results

### 2.1 Graph eco-evolutionary model

We establish an individual-based model (IBM) where individuals are structured over a trait space and a graph representing a landscape. Individuals die, reproduce, mutate and migrate in a stochastic fashion, which together results in macroscopic properties. Our formalisation results in an analytical description of the dynamics at the population level, which links emergent properties to the elementary stochastic processes that generate them.

#### 2.1.1 Microscopic description

The trait space 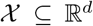 is continuous and can be split into a neutral trait space 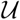 and an adaptive trait space 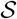. We refer to neutral traits 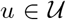 as traits that are not under selection, in contrast to adaptive traits 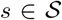, which experience selection. The graph denoted by *G* is composed of a set of vertices {*v*_1_, *V*_2_,…, *V_M_*} that correspond to demes, and a set of edges that constrain the movement of individuals across the demes. By defining 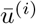 as the mean trait on *V_i_*, we quantify neutral differentiation as the variance of 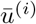 across demes and adaptive differentiation as the variance of 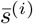 across the demes, denoted by *β_u_* and *β_s_* respectively, so that 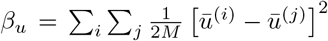. This notation is motivated by the fact that one can regard the level of differentiation as the population *β*-diversity ([78] and Supplementary Methods).

Following the Gillespie update rule [79], an individual in a deme represented by *V_i_* with trait 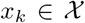 is randomly selected to give birth at rate *b*_i_(*x_k_*) and die at rate *d*(*N*^(*i*)^) = ^*N*(*i*)/*K*^, where *N*^(*i*)^ is the number of individuals on *V_i_* and *K* is the local carrying capacity. The definition of *d* therefore assumes that competition is proportional to the number of individuals in a deme, and does not depend on the individuals’ traits. Individuals reproduce asexually and the offspring resulting from a birth event inherits the parental traits, which can independently be affected by mutations with probability *μ*. A mutated trait differs from the parental trait by a random change that follows a normal distribution with variance 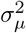 (corresponding to the continuum of alleles model [62]). The offspring can further migrate to neighbouring demes by executing a simple random walk on *G* with probability *m*. Under no selection pressure (setting (1)), individuals are only characterised by neutral traits so that 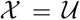. For an individual in a deme with trait *x_k_* ≡ *u_k_* we define *b_i_*(*x_k_*) ≡ *b*, so that the birth rate is constant. This ensures that neutral traits do not provide any selective advantage. Under heterogeneous selection pressure (setting (2)), the vertices of the graph are further labelled by a set of habitats ***θ*** = {*θ*_1_, *θ*_2_,…, *θ_M_*} that specify the optimal adaptive trait value *θ_i_* on *V_i_*. It follows that, for an individual with traits 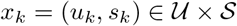, we define

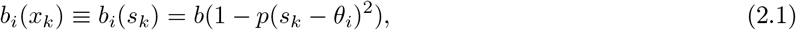

where *p* is the selection strength [74]. This ensures that on *V_i_* the maximum birth rate on *V_i_* is *s_k_* = *θ_i_*, which results in a differential advantage that acts as an evolutionary stabilising force. For the sake of the study, we assume that habitats are binary and symmetric, so that 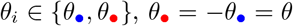, where *θ* can be viewed as the habitat heterogeneity [74].

#### 2.1.2 From stochastic dynamics to emergent deterministic properties

The model can be formulated as a measure-valued point process ([59] and Supplementary Methods). Under this formalism, we demonstrate in Supplementary Methods how the population size and trait dynamics show a deterministic behaviour when a stabilising force dominates. As a byproduct, the dynamics of the macroscopic properties can be readily expressed with deterministic differential equations, connecting emergent patterns to the processes that generate them. In particular, under the setting of no selection pressure competition stabilises the population size, and the dynamics can be considered as deterministic and expressed as

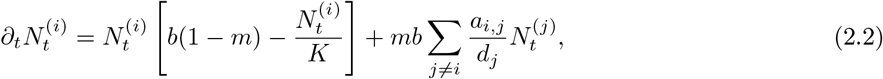

where 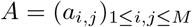 is the adjacency matrix of the graph *G* and *D* = (*d*_1_, *d*_2_,…, *d_M_*) is a vector containing the degree of each vertex. The first term on the right-hand side corresponds to a logistic growth, which accounts for birth and death events of non-migrating individuals. The second term captures the gains due to migrations, which depend on the graph topology. Assuming that all vertices with the same degree have an equivalent position on the graph, consisting in a “mean field” approach (see Methods), one can obtain a closed-form solution from Eq. (2.2), which shows that local population size *N*^(*i*)^ scales with 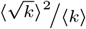, where 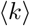 is the average vertex degree and 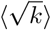 is the average square-rooted vertex degree. The quantity 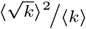, that we further denote by *h_d_*, relates to the heterogeneity in vertex degree and can therefore be viewed as a measure of heterogeneity in connectivity [80]. Complementary numerical simulations illustrate that *h_d_* can explain differences in population size for complex graph topologies with varying migration regimes (Fig. 2.2A). This analytical result connects with theoretical work on reaction diffusion processes [80] and highlights that irregular graphs result in unbalanced migration fluxes, which affect the ecological balance between births and deaths. Highly connected demes present an oversaturated carrying capacity (*N*^(*i*)^ > *K*, see Methods), increasing local competition and lowering total population size compared with regular graphs (Fig. 2.2A). Because populations with small sizes experience more drift ([67], Fig. S1B), this result suggests that graph topology not only affects neutral differentiation through population isolation, but also has a non-trivial effect on neutral differentiation by affecting population size.

**Figure 2.1:**
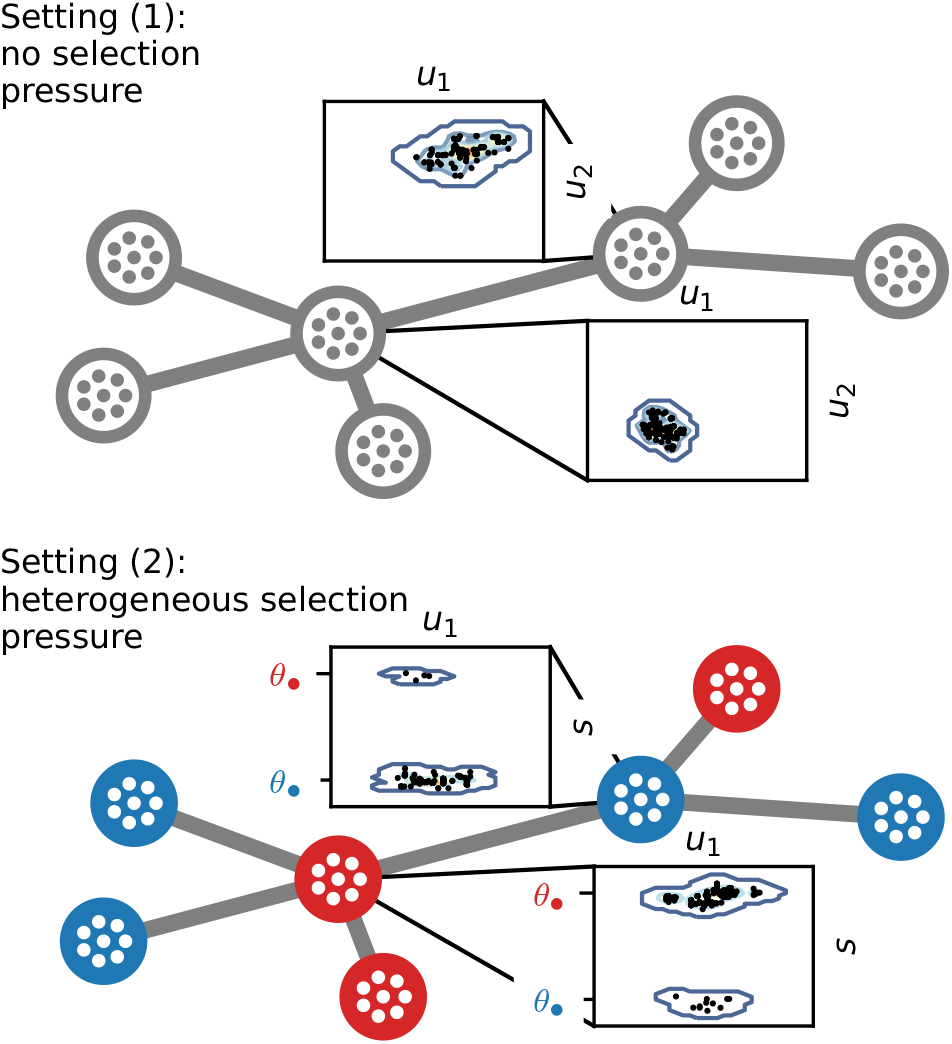
Graphical representation of the population structure. The upper panel corresponds to setting (1) with no selection pressure. In this case, individuals are characterised by a set of neutral traits 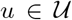. The scatter plots represent a projection of the first two components of *u* for local populations at time *t* = 500, obtained from a simulation. The lower panel corresponds to setting (2) with heterogeneous selection pressure. In this setting, individuals are additionally characterised by adaptive traits 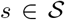. Blue vertices favour the adaptive trait optimum 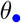, while red vertices favour 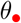. The scatter plots represent a projection of the first component of *u* and *s* for local populations at time *t* = 500, obtained from a simulation. The majority of individuals are locally adapted and have an adaptive trait centred around the trait optimum, but some less adapted individuals originating from neighbouring vertices are also present. *m* = 0.1.

**Figure 2.2:**
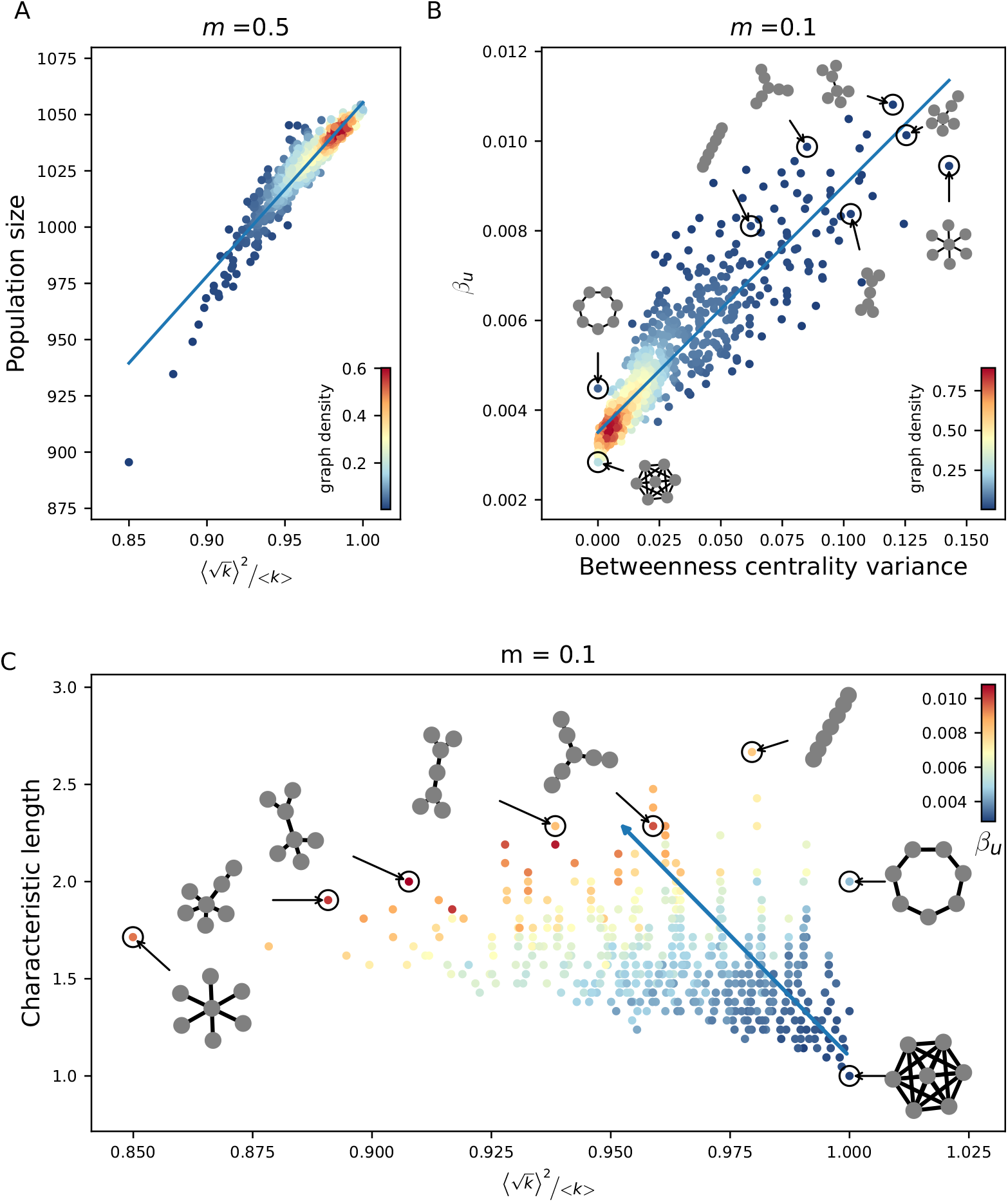
Effect of topology metrics on neutral differentiation *β_u_* and population size under no selection pressure (setting (1)). Each plot represents results from simulations of the individual-based model (IBM) on all undirected connected graphs with seven vertices. (A) Response of neutral differentiation *β_u_* to betweenness centrality variance for *m* = 0.1. This topology metric best correlates with *β_u_* for varying migration regimes (Table S1 and Fig. S5). (B) Response of population size *N* to heterogeneity in vertex degree 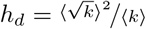, for *m* = 0.5. (B) shows that insights obtained from the mean field approximation (see Methods, Eq. (5.2)) hold for more complex graphs and varying migration regimes. In (A) and (B) the colour scale corresponds to the proportion of the graphs with similar *x* and *y* axis values (graph density), while the blue line corresponds to a linear fit. (C) Response of *β_u_* to *h_d_* and characteristic length. These two topology metrics appear in the best linear model with two predictor variables describing the effect of graph topology on *β_u_* (Table S3). In (C), the colour scale corresponds to the average *β_u_* observed for the graphs with similar *h_d_* and characteristic length values, while the blue arrow represents the gradient ∇*f* where *f* (*h_d_*, characteristic length) = *β_u_* corresponds to the linear model.

Nonetheless, the stochasticity of the processes at the individual level can propagate to the macroscopic level and significantly affect the evolutionary trajectory of the populations. In particular, the mean value of the populations’ neutral trait evolves in a stochastic fashion because it is not influenced by a stabilising force, as neutral traits do not provide any selective advantage (see Supplementary Methods and Fig. S3). Those fluctuations complicate an analytical underpinning of the dynamics, and in this case numerical simulations of the IBM offer the most effective approach to evaluate the emergent properties.

### 2.2 Effect of graph topology on neutral differentiation under no selection pressure

We study a setting with no selection pressure (setting (1)), and investigate the effect of the graph topology on neutral differentiation. When migration is limited, individuals’ traits within demes are coherent but stochastic drift at the population level generates neutral differentiation across the demes [81]. Migration attenuates neutral differentiation because it has a correlative effect on local trait distributions, and we expect that the intensity of the correlative effect depends on the topology of the graph. We consider various graphs with identical number of vertices and run simulations of the IBM to obtain the neutral differentiation level attained at a time long enough for transient population dynamics to vanish. We then examine how topology metrics summarising the graph properties explain the discrepancies in neutral differentiation *β_u_* across the simulations.

We compute the 30 scalar graph metrics listed in [77] and assess their Pearson correlations with *β_u_* for varying migration rate *m* (Fig. S5 and Table S1). We include mean population size and the heterogeneity in degree *h_d_* (see previous section) within the list of metrics, to disentangle whether heterogeneity in connectivity only affects neutral differentiation by reducing population size, or if it has more profound consequences for neutral differentiation. Although such correlations depend on the migration regime (Table S1 and Fig. S8), we find that betweenness centrality variance and characteristic length yield the strongest correlations overall (*ρ* = 0.89 and *ρ* = 0.87, respectively; Fig. 2.2B and Table S1), while population size shows a lower correlation *(ρ =* 0.53). Betweenness centrality variance measures heterogeneity in vertex centrality [77] and characterises the graph heterogeneity in connectivity. We further find that related metrics (eigenvector centrality variance, edge betweenness centrality variance, *h_d_*; Fig. S5) also correlate with *β_u_* (|*ρ*| > 0.7; Table S1). On the other hand, characteristic length measures the average vertex centrality and therefore quantifies the graph connectivity. We observe that related metrics (edge betweenness centrality mean, eigenvector centrality mean, closeness centrality mean, edge density and algebraic connectivity; Fig. S5) correlate with *β_u_* (|*ρ*| > 0.7; Table S1).

We then perform a linear regression analysis with two predictor variables for different migration regimes in order to evaluate the concurrent effect of the topology metrics on *β_u_*. We identify metrics quantifying graph connectivity and heterogeneity in connectivity as the best combination of explanatory variables (Table S3 and Fig. S5). The best linear model involves the characteristic length of the graph and the heterogeneity in degree *h_d_*, with similar contributions to neutral differentiation (*R*^2^ = 0.88; Fig. 2.2C). This result is consistent with insights from the graph metapopulation model of [31], which also proposes characteristic length as a good predictor of neutral *β* diversity. Nonetheless, in contrast to the authors conclusion that the star graph presents low differentiation due to its low characteristic length, we find that the star graph supports higher differentiation than most graphs because of competitive interactions (Fig. 2.2B). Our model assumes logistic growth, and unbalanced migration fluxes lead central vertices to host more individuals than allowed by their carrying capacity. This causes increased competition that results in a higher mortality rate, so that migrants have a lower probability of further spreading their trait. Highly connected vertices therefore behave as bottlenecks, increasing the isolation of peripheral vertices [28]. We conclude that vertices with a high degree in irregular graphs reduce effective migration and consequently amplify neutral differentiation. In the absence of selection pressure, graphs with high characteristic length and high heterogeneity in degree, or similarly graphs with low connectivity and high heterogeneity in connectivity, show high levels of neutral differentiation.

### 2.3 Effect of complex spatial habitat distribution on adaptive differentiation

We next consider heterogeneous selection pressure and investigate the response of adaptive differentiation to complex spatial habitat distribution. Adaptive differentiation emerges from the adaptation to local habitat conditions, but migration destabilises adaptation because it brings maladapted migrants. We expect that differences in connectivity across habitats, captured by the spatial habitat distribution ***θ*** = {*θ*_1_, *θ*_2_,…, *θ_M_*} and the graph topology, influence the proportion of maladapted migrants and therefore affect the level of adaptive differentiation. Selection pressure, captured by the dependence of the birth rate on θ¿, stabilises the adaptive trait distribution so that it can be approximated deterministically. We demonstrate under mean field assumption how a graph can be reduced to an equivalent two-deme model. From this simplification arises a quantity, coined the habitat assortativity rθ, that further determines adaptive differentiation for complex topologies and spatial habitat distributions.

#### 2.3.1 Insights under the mean field assumption

Due to the stabilising force of selection, the stochastic fluctuations of the adaptive trait distribution are small and one can approximate the number of individuals on *V_i_* with traits 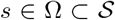 by the quantity 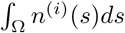, where *n*^(*i*)^ is a continuous function solution of the Partial Differential Equation (PDE)

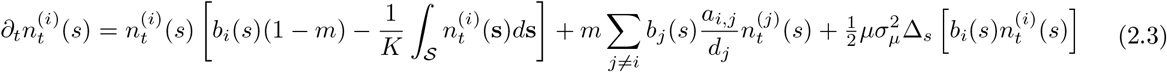

(see Supplementary Methods, and Figs. S4 and S7). Equation (2.3) is similar to Eq. (2.2), except that it accounts for selection through the dependence of the birth rate on the adaptive trait and that it incorporates a last term corresponding to mutation processes. In the case where *m* = 0, [82, 83] show that Eq. (2.3) admits a stationary solution *n*^(*i*)^ that is Gaussian with mean 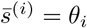, and so that 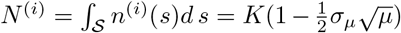. Hence, under low *μ* and *σ*, population size is similar to that in setting (1). It further indicates that, in contrast to *β_u_, β_s_* does not depend on population size, but is rather governed by Var(***θ***), the variance of the habitat distribution.

We now show under mean field assumption how Eq. (2.3) can be reduced in the general case where *m* ≥ 0 to an equivalent two-deme model. The mean field approach slightly differs from setting (1) because vertices are labelled with *θ_i_*. In this case, we assume that vertices with similar habitats have an equivalent position on the graph (see Fig. S10 for a graphical representation), so that all vertices with habitat 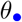 are characterised by the same adaptive trait distribution that we denote by 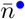. Let 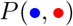 denote the proportion of edges connecting a vertex with habitat of type 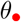 to a vertex with habitat of type 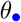, and let 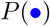 denote the proportion of vertices with habitat 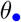. By additionally assuming that habitats are homogeneously distributed so that 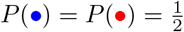, Eq. (2.3) transforms into

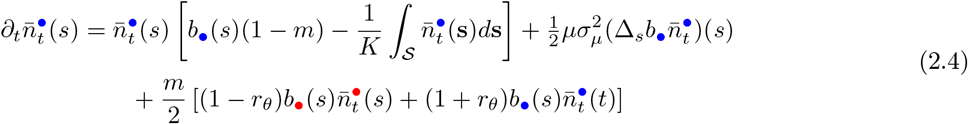

(see Methods), where we define

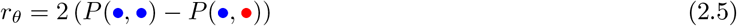

as the habitat assortativity of the graph, which ranges from −1 to 1. The case where *r_θ_* = −1 corresponds to the case where all edges map dissimilar habitats (disassortative graph), while as *r_θ_* → 1 the graph is composed of two clusters of vertices with homogeneous habitats (assortative graph). We numerically solve Eq. (2.4) and show in Fig. 2.3 how both *m* and *r_θ_* influence population size and *β_s_*. As expected, migration destabilises local adaptation and decreases population size. We find a critical threshold *m*\* that increases with *r_θ_*; when *m* > *m*\*, local adaptation cannot be sustained anymore and *β_s_* consequently vanishes (Fig. 2.3A, red curve). Figure 2.3A shows that increasing assortativity *r_θ_* systematically increases *β_s_* at an equivalent migration regime. Our analytical reduction thereby demonstrates that, under the mean field assumption, assortative graphs present high levels of adaptive differentiation. On the other hand, population size and *β_s_* rapidly decline with increasing migration on disassortative graphs, until *β_s_* vanishes completely when *m* > *m*\*.

**Figure 2.3:**
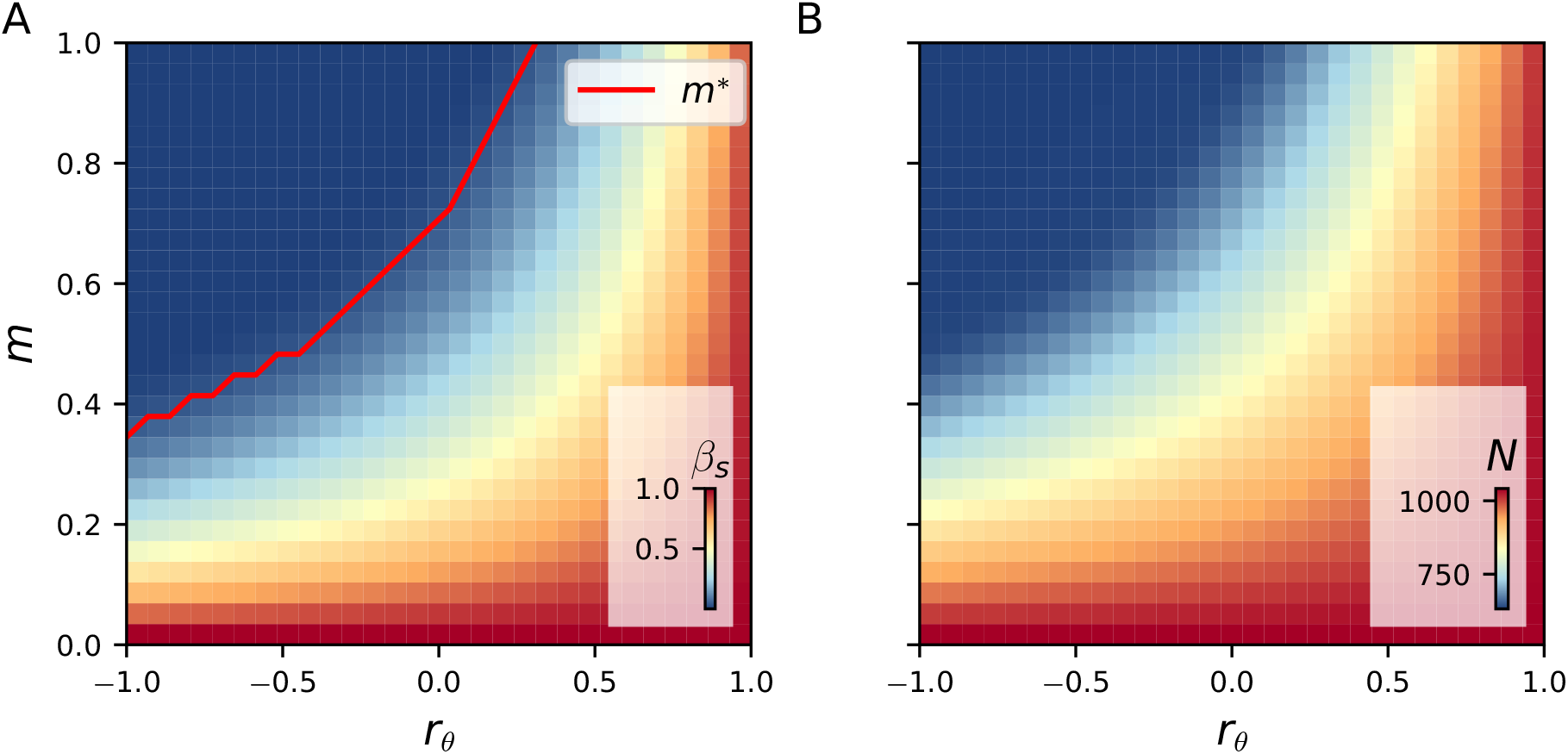
Effect of habitat assortativity *r_θ_* and migration on adaptive differentiation *β_s_* and population size under the mean field, deterministic approximation Eq. (2.4). (A) Effect of *r_θ_* on *β_s_*. The red line indicates the critical migration threshold *m*\* for which *β_s_* vanishes when *m* > *m*\*. (B) Effect of *r_θ_* on population size. (A) and (B) show that increasing *r_θ_* systematically increases *β_s_* and population size, irrespective of the migration regime.

#### 2.3.2 Relaxing the mean field assumption

In order to verify the conclusions obtained with the mean field, deterministic approximation Eq. (2.4), we generate randomly different habitat distributions ***θ*** = {*θ*_1_, *θ*_2_,…, *θ_M_*} for varying graph topology, and we compare simulations of the IBM (see Methods) with results from the PDE model Eq. (2.4) under different migration regimes (Fig. 2.4). For each combination of ***θ*** and graph, we compute the habitat assortativity *r_θ_*, using the fact that *r_θ_* can be generalised from Eq. (2.5) to any graph topology following the original definition of [84]

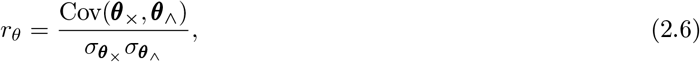

where ***θ***_×_ and ***θ***_∧_ denote the sets of habitats found at the toe and tip of each directed vertex of graph *V*, and 〈***θ***_×_〉, 〈***θ***_∧_〉 and 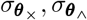 denote their respective means and standard deviations (see Supplementary Methods). Figure 2.4A,B confirm that the mean field, deterministic approximation Eq. (2.4) captures the response of *β_s_* to *r_θ_* for more general graph ensembles. On the other hand, we assess the correlation between *β_s_* and topology metrics and perform a linear regression analysis to show that the topology metrics have a negligible effect on *β_s_* (Tables S2 and S4).

**Figure 2.4:**
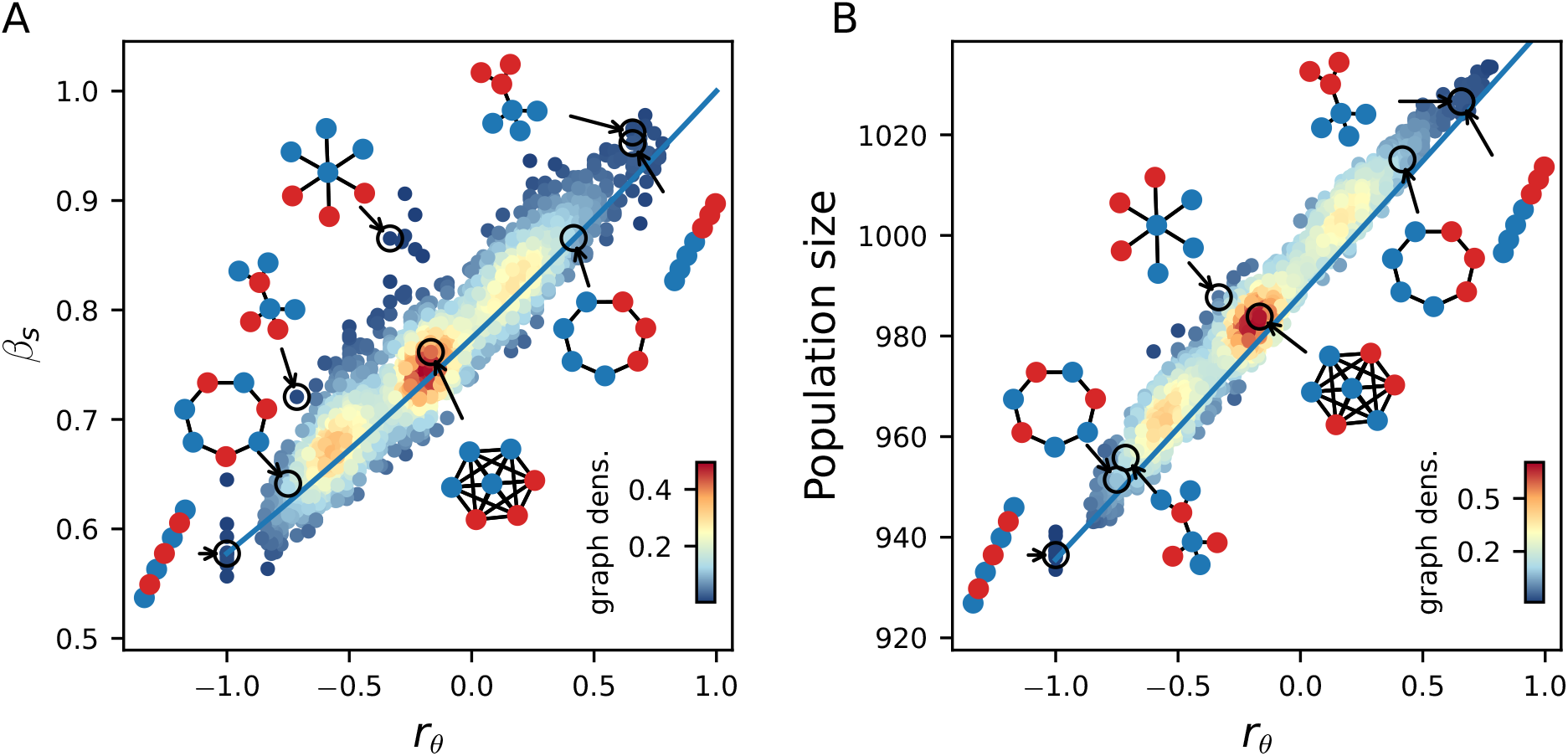
Effect of habitat heterogeneity *r_θ_* on *β_s_* and population size for general graph ensembles. Each plot represents results from simulations of the IBM on all undirected connected graphs with seven vertices and varying *r_θ_*, for *m* = 0.1. The colour scale corresponds to the proportion of the graph with similar *x* and *y* axis values (graph density). The blue lines correspond to results obtained from the mean field approximation Eq. (2.4). (A) and (B) illustrate that insights from the mean field, deterministic approximation Eq. (2.4) remain valid in the stochastic setting for complex habitat connectivity patterns.

Under strong selection, stochastic drift vanishes [85] and we find that the effect of complex connectivity patterns across habitat can be reduced to a simplified two-deme model where migration rates are altered by a factor that involves the habitat assortativity *r_θ_*. Mirrahimi et al. [83, 74] investigated a deterministic two-deme model corresponding to the IBM approximation Eq. (2.4) in the special case where *r_θ_* = −1, with the difference that they assumed that individuals continuously migrate and mutate (no dependence of the migration and mutation terms on the birth rate *b*). The authors provided a closed form for *m*\*, yielding *m*\* = *pθ*^2^, that indicates that as selection or habitat heterogeneity *θ* increases, so does *m*\* and local adaptation is sustained under higher migration regimes. On the other hand, considering the stochastic dynamics of the IBM, as selection pressure decreases the birth rate *b* provides only a marginal advantage to adapted individuals, so that stochastic drift in the populations’ mean adaptive trait rises up. In this case the role of habitat assortativity in determining the level of differentiation should be supplanted by the graph topology, so that characteristic length and heterogeneity in degree become the dominant drivers of differentiation patterns. Irrespective of the graph connectivity *per se*, we thus find that under strong enough selection the adaptive differentiation is mainly driven by the habitat assortativity *r_θ_* and vanishes for *m* > *m*\*, where *m*\* depends on the habitat assortativity of the graph *r_θ_*, the habitat heterogeneity *θ* and the selection pressure *p*.

### 2.4 Ambiguous effect of habitat assortativity on neutral differentiation

We eventually consider individuals carrying both neutral and adaptive traits (setting (2)). Under heterogeneous habitats, selection pressure promotes neutral differentiation by reducing the reproductive rate of maladapted migrants, reinforcing local population isolation [29]. We have shown that adaptive differentiation is driven by habitat assortativity, so we expect habitat heterogeneity, together with the topological metrics found in setting (1), to influence the level of neutral differentiation. We first investigate how the response of neutral differentiation to migration compares between setting (1) and setting (2) for graphs with an identical topology.We then examine how the response compares between graphs with an identical topology but different habitat assortativity. We finally consider simulations on different graphs with varying habitat assortativity to assess the concurrent effect of assortativity and topology metrics on neutral differentiation.

Migration has a fitness cost because maladapted migrants present lower fitness [71]. Under an equivalent migration regime, migrants therefore have a lower probability of reproduction, increasing the populations’ isolation compared with a scenario without selection pressure [29]. In Figure 2.5A we present simulation results obtained by varying *m* on the complete graph. This confirms that selection pressure in heterogeneous habitats reinforces neutral differentiation compared with a scenario without selection pressure. Nonetheless, previous results show that adaptive differentiation vanishes with a disassortative graph when *m* > *m*\*, implying that individuals become equally fit in all habitats. In this case, the isolation effect of heterogeneous selection pressure is lost, and Fig. 2.5A shows that *β_u_* levels in setting (2) reach the same level as in setting (1) for *m* > *m*\*. This suggests that habitat assortativity reinforces *β_u_*, as assortative graphs sustain higher levels of adaptive differentiation (Fig. 2.4). Simulations on the chain graph with varying spatial habitat distribution give support to this reasoning for high migration regimes, but show that neutral differentiation is higher for low habitat assortativity under low migration regimes (Fig. 2.5B). Assortative graphs are composed of large clusters of vertices with similar habitats, within which migrants can circulate without fitness losses. Local neutral trait distributions become more correlated within those clusters, resulting in a decline in neutral differentiation for assortative graphs compared with disassortative graphs. Figure 2.5B therefore highlights the ambiguous effect of habitat assortativity *r_θ_* on neutral differentiation. *r_θ_* reinforces neutral differentiation by favouring adaptive differentiation, but at the same time it decreases neutral differentiation by decreasing population isolation within similar habitat clusters. A criterion for determining when *r_θ_* favours or depresses neutral differentiation should depend on habitat heterogeneity *θ* and selection pressure p, but is not trivial and remains to be determined. We finally compare the effect of habitat assortativity on neutral differentiation to the effect of the connectivity metrics found in setting (1). We perform a linear regression analysis with three predictor variables on simulation results obtained for different graphs with varying habitat distribution, using as predictors *r_θ_*, characteristic length and heterogeneity in degree *h_d_* (Fig. 2.5B,C and Table S5). The linear model explains the discrepancies in neutral differentiation across the simulations for varying migration regimes (*R*^2^ > 0.83; Table S3), and we find that both *r_θ_* and the two topology metrics contribute similarly to neutral differentiation. This analysis suggests that the effects of habitat assortativity and connectivity of the graph add up under heterogeneous selection pressure. A change in sign of the coefficient for *r_θ_* appears when comparing models fitted for low and high migration regimes, verifying that the opposite effect of habitat assortativity on *β_u_* found on the chain graph holds for general graph ensembles. Characteristic length and heterogeneity in degree therefore drive neutral differentiation with and without heterogeneous selection pressure (setting (1) vs setting (2)). Habitat assortativity *r_θ_* becomes an additional driver of neutral differentiation under heterogeneous selection pressure. In contrast to the non-ambiguous, positive effect of habitat assortativity on adaptive differentiation, *r_θ_* can amplify or depress neutral differentiation depending on the migration regime considered.

**Figure 2.5:**
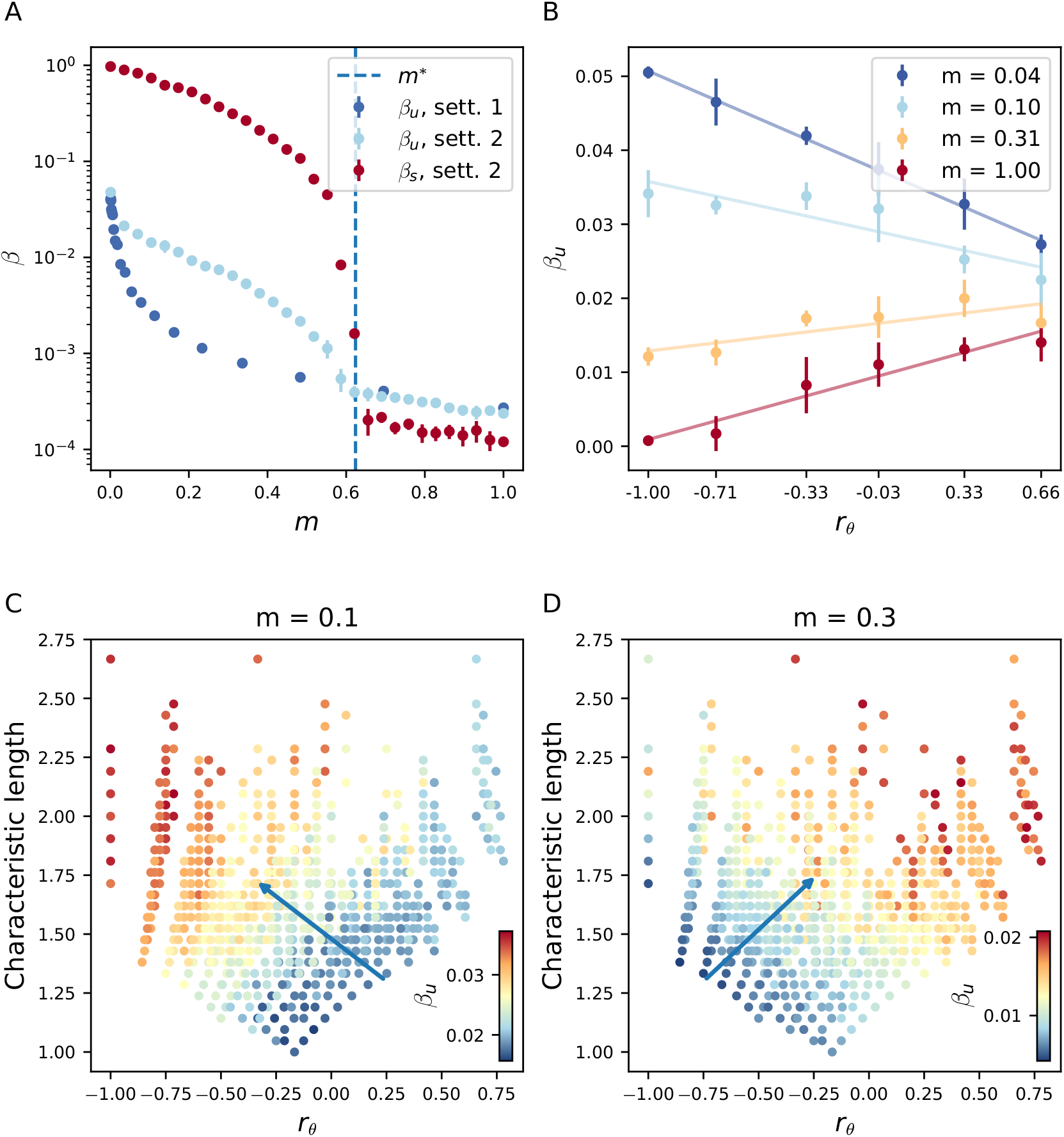
Effect of habitat assortativity *r_θ_* and connectivity metrics on neutral differentiation *β_u_* under heterogeneous selection pressure (setting (2)). (A) Comparison of the response of *β_u_* to migration in setting (2) to the response of *β_u_* in setting (1) (no selection) for the complete graph. (A) shows that *β_u_* is higher in setting (2) than in setting (1) when *m* < *m*\*. However, adaptive differentiation *β_s_* is lost for *m* > *m*\*, and in this case the *β_u_* level in setting (2) reaches the level of *β_u_* in setting (1). The dashed blue line corresponds to the critical migration regime *m*\* predicted by the mean field, deterministic approximation Eq. (2.4). (B) Response of *β_u_* to *r_θ_* and migration for the chain graph. (B) illustrates the ambiguous effect of *r_θ_* on *β_u_* and shows that under identical topology, *r_θ_* correlates positively with *β_u_* for high migration regimes, but correlates negatively for low migration regimes. (C) and (D) Concurrent effect of *r_θ_* and characteristic length on *β_u_* for low and high migration regimes. (C) and (D) show results from simulations of the IBM on all undirected connected graphs with seven vertices and varying *r_θ_* (see Methods). (C) and (D) illustrate the additive effects of *r_θ_* and connectivity metrics on *β_u_*, and confirm that the ambiguous effect of *r_θ_* found for the chain graph holds for general graph ensembles. In (C) and (D) the blue arrows represent the gradient ∇*f*, where *f*(*r_θ_*, characteristic length, *h_d_*) = *β_u_* corresponds to fitted linear model with three predictor variables.

## 3 Discussion

Using analytical tools and simulations, we have built upon an individual-based model and a graph representation of landscapes to investigate how landscape features drive diversification. Characteristic length, heterogeneity in degree and habitat assortativity are the main drivers of differentiation with respect to our modelling assumptions, and we have demonstrated how those properties relate to the underlying mechanisms [86, 87]. Our results connect with correlative studies that have already associated species richness with a variety of metrics used as surrogates for connectivity [7, 88, 5, 89], connectivity heterogeneity [10, 9, 11, 90] and habitat heterogeneity [91, 6, 92]. Techniques to project real landscapes on graphs have been proposed in [35, 93], and we illustrate their applicability in Fig. S12, that demonstrates how to characterise large geographical areas by directly using graph-based topology metrics. This approach could eventually be used in empirical studies to gain further understanding on the origin of spatial biodiversity patterns. Overall, our results propose topology metrics that can support empiricists to better connect spatial biodiversity patterns to generating eco-evolutionary mechanisms and landscape features.

In the absence of selection pressure, neutral differentiation is more pronounced in graphs with high characteristic length, but is also associated with heterogeneity in degree (Fig. 2.2C). Characteristic length can be viewed as a measure of spatial dimensionality [35], and this result is consistent with empirical studies [7, 88, 5, 89], classical theoretical works based on the stepping stone model [62, 65, 66], as well as the graph metapopulation model of Economo et al. [31]. However, these works did not consider ecological interactions between individuals within populations, and our results show that such interactions can substantially influence the effect of graph topology on neutral differentiation. Our model assumes that population growth is limited by the local carrying capacity, which becomes saturated on highly connected vertices in irregular graphs. As a consequence, these vertices behave as bottlenecks and amplify the isolation of peripheral vertices [28], so that heterogeneity in connectivity appears as an equally strong driver of neutral differentiation in graphs (Fig. 2.2B,C). This behaviour should be prevalent in patchy landscapes where interspecific competition is high due to limiting resources [30]. While it has been acknowledged that reaction diffusion processes behave very differently in graphs with heterogeneous connectivity compared with homogeneous graphs [80, 94, 95], we further document how competitive interactions, as reaction processes, can influence the emergence of phenotypic differentiation in graphs. The role of heterogeneity in connectivity cannot be captured with classical metapopulation and quantitative genetics models [96] or with models of evolutionary dynamics in graphs [47], as they assume constant population size. Our study highlights that heterogeneity in connectivity can reinforce differentiation patterns through the creation of unbalanced migration fluxes which affect ecological equilibrium.

Habitat assortativity *r_θ_* is a useful indicator for assessing how complex spatial habitat distributions modulate local adaptation and adaptive differentiation [97, 98]. While adaptation has been extensively studied along environmental clines [81, 64, 99, 100, 71, 101, 68], landscapes can be patchy and assuming regularity might be unrealistic [35]. Graphs can capture irregularity in connectivity across habitats [102, 93], and our results indicate that, irrespective of the graph topology, adaptive differentiation is mainly driven by the habitat assortativity *r_θ_* (Fig. 2.4), which can be regarded as the habitat spatial auto-correlation [84]. Habitat assortativity *r_θ_* can be obtained for general habitat distributions that might not be neither binary nor symmetric (see [103, 84] and Fig. S12), and we expect our conclusions to hold in such cases. Low habitat assortativity reinforces the “swamping” effect of gene flow proposed by Haldane [100, 97] and limits local adaptation, so that *r_θ_* can be viewed as the spatial scale of habitat variation that conditions local adaptation [104, 105]. Montane regions can support extremely varied habitats within a small spatial scale (see [8] and Fig. S12), and in this case habitat assortativity should be low. This implies that for taxa to be locally adapted, a low migration rate is required, which might explain why many species in such environments have small ranges [106, 107]. Our results predict that under an equivalent migration regime, populations structured over assortative habitats are larger, support higher adaptive differentiation, and can be locally adapted even in the case where migration rates are strong.

Spatial eco-evolutionary feedbacks in heterogeneous habitats critically affect diversification [108, 109, 69]. While most eco-evolutionary studies have investigated diversification by considering a unique adaptive trait [68, 99, 100, 71, 101, 110], distinguishing between neutral and adaptive differentiation is crucial [19] and our work underlines their distinct responses to landscape properties (Fig. 2.3A vs. Fig. 2.5C). Our study builds upon recent mathematical models that consider the co-evolution of neutral and adaptive traits [75, 111, 76] and extends those works to a spatial context. Our work provides an analytical framework to the concept of isolation by environment (IBE) [28], which has been suggested to be one of the most important mechanisms governing differentiation in nature [29]. Heterogeneous selection pressure leads to more isolation by modifying the fitness of migrants [71], and therefore affects the level of neutral differentiation (Fig. 2.5A) [30]. In empirical studies an ambiguous response of species richness to habitat heterogeneity has been observed [112, 91, 92]. Our work reconciles this apparent inconsistency and proposes a mechanism by which habitat assortativity, relative to the migration regime, controls the direction of the effect of habitat heterogeneity on diversification (Fig. 2.5C,D). Patchy, heterogeneous habitats can promote neutral differentiation as a result of selection pressure that reduces effective migration [92]. Nonetheless, adaptive differentiation decreases significantly when migration is high. In this case, neutral differentiation should be higher in landscapes with more aggregated habitats [104]. Our study suggests that habitat assortativity should be considered a unifying metric for understanding the causal link between habitat heterogeneity and species richness in complex environments [92].

## 4 Conclusion

From a rigorous analytical description of micro-evolutionary processes explicitly accounting for spatial dynamics over graphs, we have established how diversification can emerge at the population level from eco-evolutionary feedbacks in complex landscapes. Our study formalises verbal propositions on how differentiation emerges on ecological time scales from the interplay between spatial dynamics, the co-evolution of neutral and adaptive traits, and landscape properties [113, 28, 29, 104]. In agreement with findings from empirical studies [7, 88, 5, 89, 91, 6, 92] and previous theoretical considerations [31, 62, 65, 66], characteristic length (which relates to landscape connectivity) and habitat assortativity (which relates to habitat spatial auto-correlation) emerge as core drivers of differentiation. We have further shown how the migration regime dictates whether neutral differentiation positively or negatively correlates with habitat heterogeneity. Additionally, our work highlights that heterogeneity in connectivity should equally be a strong driver of differentiation because highly connected demes behave as bottlenecks, increasing the isolation of peripheral demes. The framework laid out here is a promising approach for studying complex adaptive systems, as it can elucidate how macroscopic properties emerge from microscopic processes [14]. It could be used in other fields such as in linguistics [114] or economics [115], where agents interact and are also structured over complex spatio-evolutionary structures.

## 5 Methods

### 5.1 Mean field approximation

In setting (1), the mean field approach involves the assumption that all vertices having the same degree are equivalent. For this, let *P*(*k, k*′) denote the proportion of edges that map a vertex with degree *k* to a vertex with degree *k*′, and consider the average population size 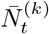 in each vertex with degree *k*. An individual has probability 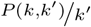 to migrate from a vertex with degree *k*′ to a vertex with degree *k*. Viewing 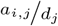 as the probability that an individual on *V_i_* chosen for migration goes to *V_j_*, Eq. (2.2) then transforms into

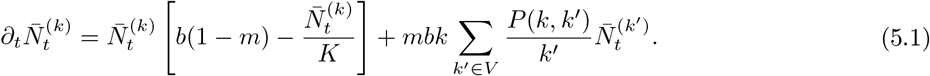

Assuming uncorrelated graphs for which 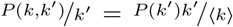, where 〈*k*〉 denotes the average degree of the graph [116, 117], yields

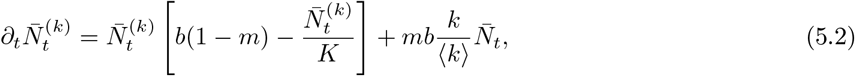

where

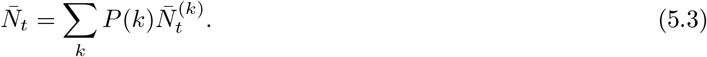

When solving for the stationary state and setting *m* = 1, one obtains 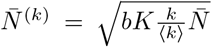 from Eq. (5.2). Combining this with Eq. (5.3) yields 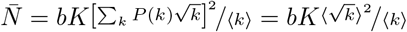.

In setting (2), the mean field approach involves the assumption that all vertices with a similar habitat are equivalent. In this case, an individual from a vertex with habitat 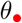 has the probability 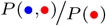 to migrate to a vertex with habitat 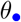, and therefore Eq. (2.3) transforms into

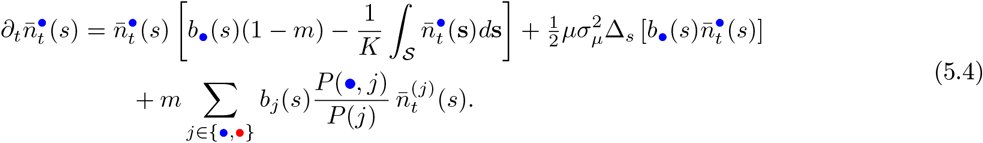

Considering that 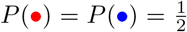 (habitats are equally distributed), 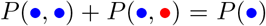 (sum of conditional expectations) and 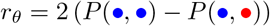 (Eq. (2.5)), one obtains

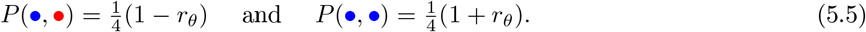

Combining Eq. (5.5) with Eq. (5.4) yields Eq. (2.4). We show in Supplementary Methods how one can derive Eq. (2.5) from the general definition of assortativity given in Eq. (2.6) and initially introduced in [84].

### 5.2 Numerical simulations

We performed Monte Carlo simulations by running five replicate simulations for each result presented, with *b* =1, total time span *t* = 500, local carrying capacity *K* = 150, selection strength *p* = 1, mutation rate *μ* = 0.1, and mutation range *σ_μ_* = 5 · 10^−2^. This parameter choice corresponds to 10^6^ birth and death events, which made it possible to obtain results in a reasonable amount of time without including transient population dynamics. In settings (1) and (2), we ran simulations on all of the 853 undirected connected graphs with 7 vertices (listed at http://oeis.org/A001349). For setting (2), we randomly generated different spatial habitat distributions and selected the ones with a unique *r_θ_* value, corresponding to a total of 2537 different simulations for fixed *m*. We then computed diversity metrics that we further averaged over the last time steps and across the replicates. Since neutral diversity dynamics are characterised by large quadratic variations, we simulated individuals with *d* = 300 neutral traits, where each trait can independently be affected by mutations. Diversity metrics were then obtained from the average *α_u_, β_u_, γ_u_* for each trait. This reduced the variance of the numerical simulations and is also biologically meaningful because populations are characterised by many traits, most of which are neutral [19]. As initial conditions, *MK* individuals were homogeneously distributed across all of the vertices, with traits centred at 0 and with standard deviation *σ_μ_*. Graph metrics used for the meta-analysis were calculated using the **LightGraphs.jl** library [118]. We numerically solved the PDEs with a finite difference scheme using **DifferentialEquations.jl** [119], ensuring that the domain was large enough to avoid border effects.

#### EvoId.jl

We have implemented the modelling framework in a computationally efficient and user-friendly package, **EvoId.jl**, written in the Julia programming language and freely accessible on GitHub. The user can specify any combination of spaces over which the populations are structured, together with birth and death functions and specific update rules. The intention is to popularise the tool and encourage the community to investigate this framework for multi-disciplinary case studies.

## Author contributions

V.B. and L.P. designed research; V.B. performed research; V.B. and L.P. wrote the paper. The authors declare no competing interest.

## Acknowledgements

We thank Thomas Poulet, Sylvian Billiard, Sepideh Mirrahimi, Heike Lischke, Joshua Payne, Conor Waldock, Flora Desmet, Benjamin Flück and Alexander Skeels for helpful discussions and comments on the manuscript. L.P. was supported by the Swiss National Science Foundation grant (N° 310030_188550).

## Data accessibility

The code used in this article is available online at https://gitlab.ethz.ch/publications/neutral-and-adaptive-diversification-in-spatial-graphs

## Supplementary Methods

### Mathematical construction of the model

The model is a measure-valued point process [59], so that individuals are represented as dirac functions 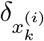, where 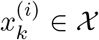 corresponds to the traits’ value of individual *k* located on vertex *V_i_*. Under this formalism, the population on *V_i_* is represented as a sum of dirac functions 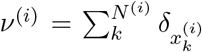, where *N*^(*i*)^ is the local population size. It follows that the time variation of the process can be described by the so-called infinitesimal generator *L*, defined for all real valued functions *ϕ* by

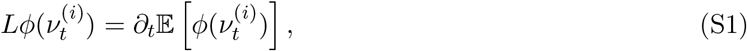

(see e.g. [120] for an introduction on infinitesimal generators). Equation (S1) provides the expected time variation at time *t* of e.g. the population size by choosing 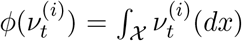. Recall that we denote by *b_i_* the birth rate on vertex *V_i_*, by *d* the death rate, by *μ* the mutation probability, by *m* the migration probability, by 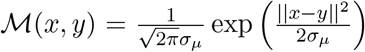 the mutation kernel, by *K* the carrying capacity, by 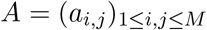 the adjacency matrix of the graph *G*, and by *D* = (*d*_1_, *d*_2_,…, *d_M_*) the vector containing the degree of each vertex. In order to explicitly write the generator *L*, let us remind ourselves that five events of different nature can alter the number of individuals with trait *x* on vertex *V_i_*:

- an individual on *V_i_* with trait *x* can give birth to an offspring with similar trait that stays on *V_i_*, at rate (1 – *μ*)(1 – *m*)*b_i_*(*x*),
- an individual on *V_i_* with trait *y* can give birth to an offspring with trait *x* that stays on *V_i_*, at rate 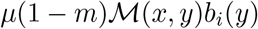,
- an individual on *V_i_* with trait *x* can die, at rate 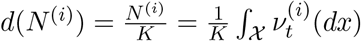,
- an individual on *V_j_* with trait *x* can give birth to an offspring that migrates to *V_i_*, at rate 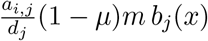,
- an individual on *V_j_* with trait *y* can give birth to an offspring that migrates to *V_i_*, at rate 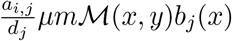.

Summing over all all individuals and all vertices yields

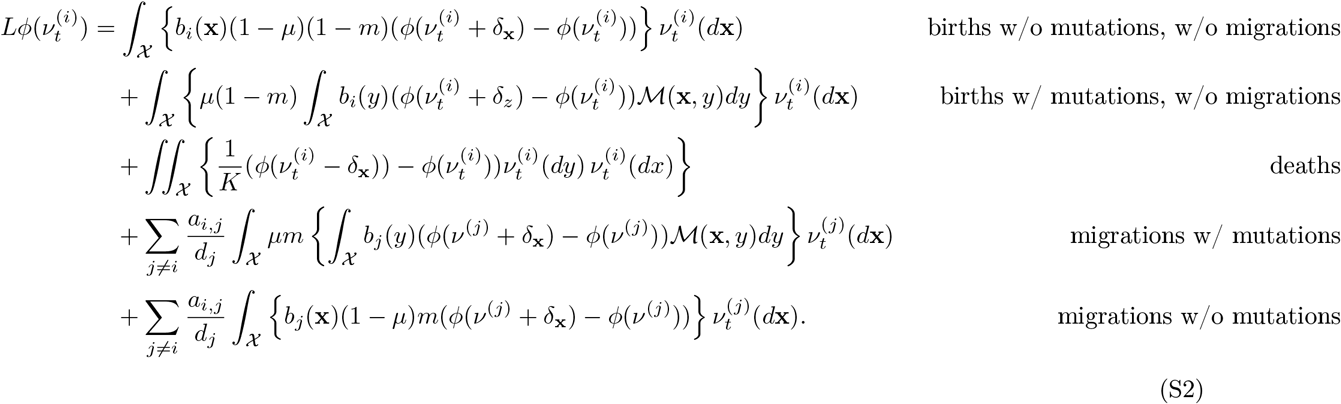

Taking expectations in Eq. (S2), one can obtain an equation for the mean trajectory of the quantity of interest, 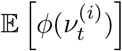. Nonetheless, Eq. (S2) involves an integral with respect to 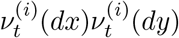, preventing one to obtain an explicit solution by doing so. It is therefore unclear whether one can gain insights on the stochastic dynamics from Eq. (S2) without simplifying assumptions, and we refer to [58] for a detailed discussion on the topic.

### Deterministic approximation

A strategy to overcome the difficulties encountered above is to assimilate the process to its mean trajectory, assuming that 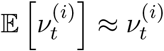 and further approximating 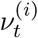 by a continuous deterministic function 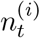. Such strategy inherently neglects the stochasticity of the process, which is reasonable provided that a force dampens the stochastic fluctuations of the quantity of interest.

### Setting 1

Consider setting (1) and recall that in this setting we define for 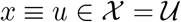

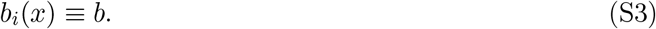

Applying the strategy mentioned above and choosing 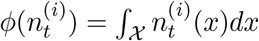, Eq. (S2) transforms into the deterministic approximation of the population size dynamics given in the main-text by

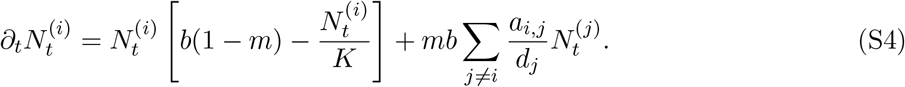

Competition stabilises the population size dynamics, which behaves deterministically. This is supported by Fig. S6A, that shows how Eq. (S4) accurately describes the population size for varying migration regimes. Nonetheless, stochastic fluctuations drive the dynamics of the neutral trait distribution. Attempting to characterise it with the same strategy by choosing this time 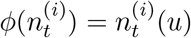 yields

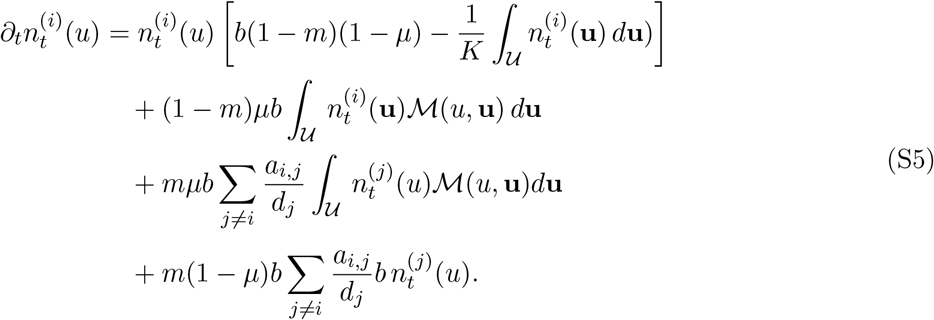

Solving for Eq. (S5), one can show that the variance of 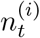 continuously grows in time (see Fig. S6) and tends to infinity as time goes to infinity, which is an unrealistic behaviour considering finite populations. Intuitively, this reflects the fact that no stabilising force acts on the neutral trait distribution, so that random fluctuations play a major role in driving the dynamics of the stochastic process. Figure S6 shows how IBM trajectories significantly differ from Eq. (S5), and Fig. S3 illustrates how diversity metrics obtained from Eq. (S5) do not match those obtained from simulations of the IBM.

### Setting (2)

Nonetheless, the adaptive distribution can successfully be approximated by a deterministic description because in contrast to the neutral trait dynamics, selection pressure acts as a stabilising force and stabilises the populations’ adaptive trait, dampening the stochastic fluctuations. Consider setting (2) and recall that in this setting we define for 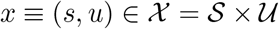

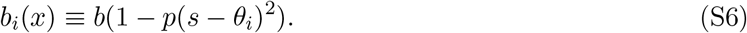

Applying the same strategy as above to characterise the adaptive trait distribution 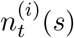 by choosing 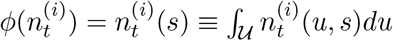, Eq. (S2) transforms into

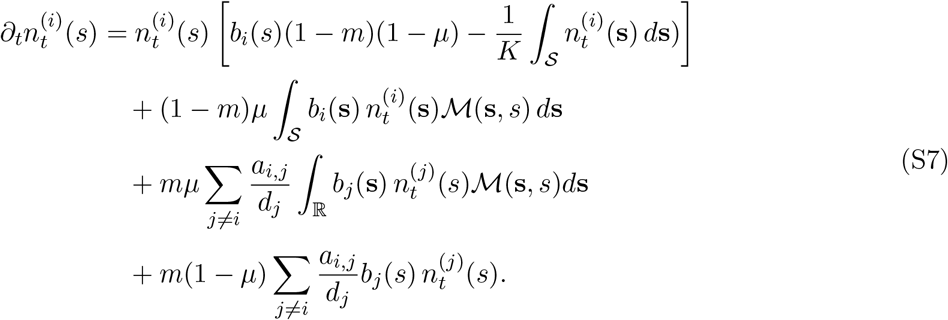

Assuming that the variance of the mutation kernel is small, one can use a diffusion approximation for the mutation term [82, 72, 74]

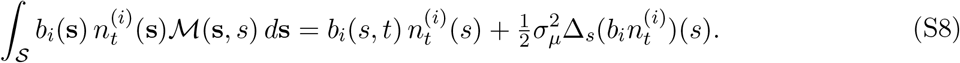

Neglecting the terms in *mμ*, we obtain

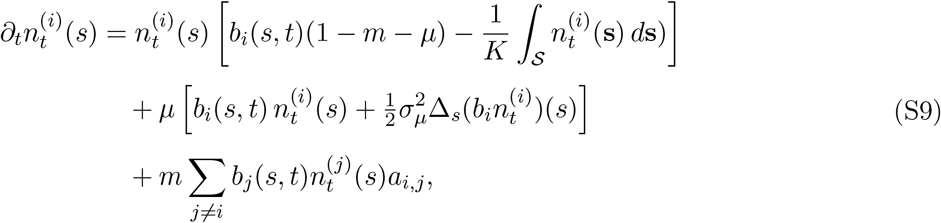

which after rearranging terms yields the elegant deterministic approximation of the adaptive trait dynamics

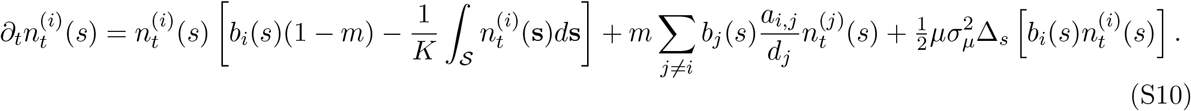

Setting *m* = 0, [74] shows that Eq. (S10) admits a stationary solution that is Gaussian, with variance 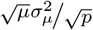. Therefore, the variance of the adaptive trait distribution stabilises to a finite value. Intuitively, this reflects the fact that the random fluctuations of the adaptive trait distribution are dampened by the stabilising force of selection. Provided that the selection strength *p* is large enough, Eq. (S10) is a good approximation of the adaptive trait distribution obtained from the stochastic process. Figure S7 shows how IBM trajectories are similar to the ones obtained from Eq. (S5), and Fig. S4 illustrates how diversity metrics obtained from Eq. (S5) match those obtained from simulations of the IBM.

### Diversity partitioning

Although we have limited ourselves to characterising *β* diversity, we introduce here the *α* and the *γ* diversity and show how those definitions allow to recover the classical additive diversity partitioning [78]. Such quantities prove particularly useful to characterise trait distributions, and are used in Figures S3 and S4. In the following we think of *x* as denoting whether the neutral or the adaptive trait, and we denote the trait mean on *V_i_* as 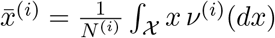.

Local *α* diversity observed on *V_i_* is defined as the variance of the local trait distribution

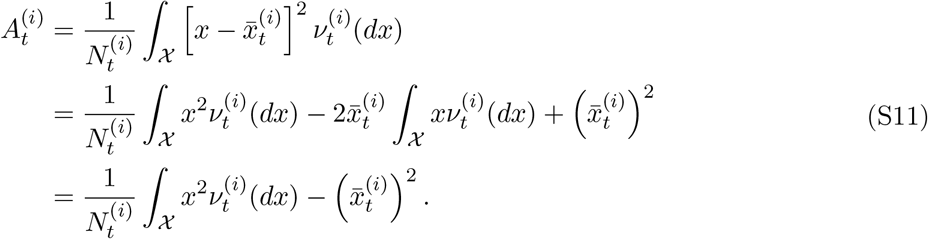

We denote the mean *α* diversity as 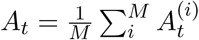 and we denote the expected mean *α* diversity as 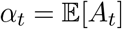.

We recall that the observed *β* diversity is defined as the variance of the local trait distribution

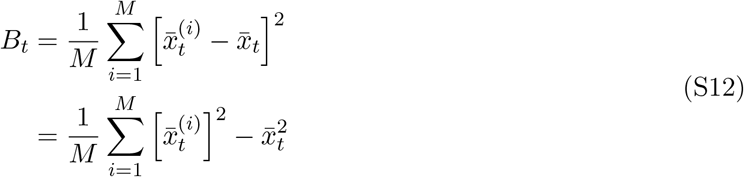

where 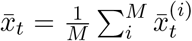 is the mean of the total trait distribution. We further denote the expected *β* diversity as 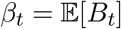.

Finally, we define *γ* diversity as the variance of the total trait distribution

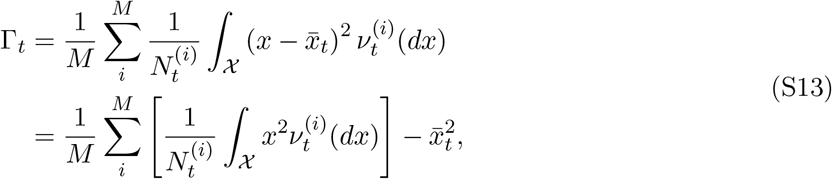

and set 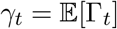.

Combining Eqs. (S11) and (S12), we obtain

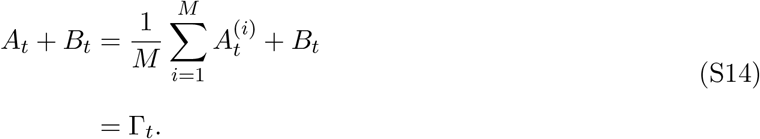

Taking expectations, we obtain

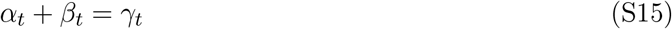

and therefore recover the classical additive diversity partitioning [78].

### Derivation of the habitat assortativity metric *r_θ_* in binary environments

We demonstrate here how the habitat assortativity *r_θ_* relates to the conditional probability of habitats being connected, and show how *r_θ_* simplifies under mean field assumption.

Following the original definition of [84], habitat assortativity *r_θ_* is defined as the Pearson correlation of environmental conditions *θ* at either ends of the vertices *V* of graph *G*, that is

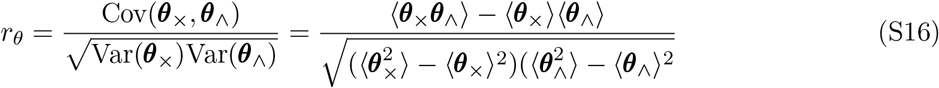

where ***θ***_×_, ***θ***_∧_ denote the sets of environmental conditions found at the toe and tip of each directed vertex of graph *V*, and 〈***θ***_×_〉, 〈***θ***_∧_〉 denote their respective mean values.

Let 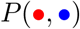 be the proportion of edges that connect a vertex with habitat 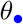 to a vertex with habitat 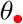. One can also view 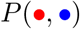 as the conditional probability that a vertex of type 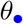 is connected to a vertex of type 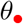. Denote by 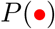 the proportion of vertices that are of type 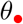. First observe that for undirected graphs, one has 〈***θ***_×_〉 = 〈***θ***_∧_〉, and 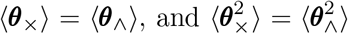. Assuming that habitats are symmetric and binary, one has that 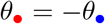. Then

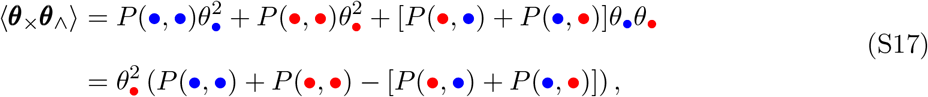

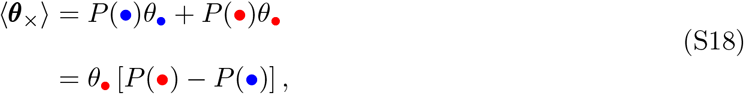

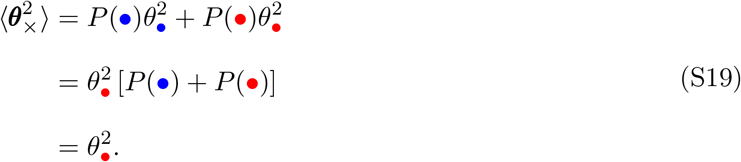

Combining Eq. (S17), Eq. (S18), and Eq. (S19) with Eq. (S16) one gets

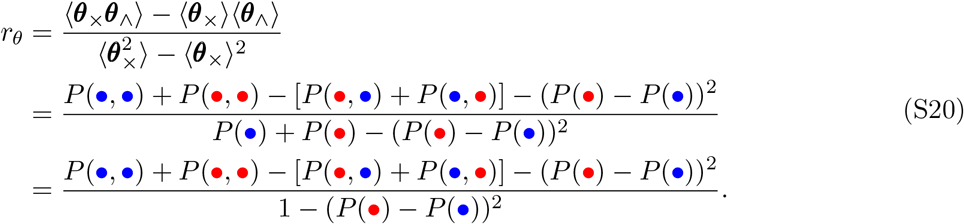

Assuming that habitats are homogeneously distributed, we have 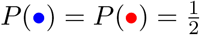 and thus we obtain

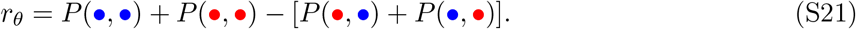

The mean field approximation involves the assumption that all vertices with similar habitats are equivalent in terms of their connections with other habitats, so that 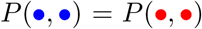 and 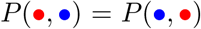, which yields 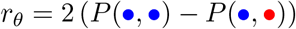.

**Figure S1:**
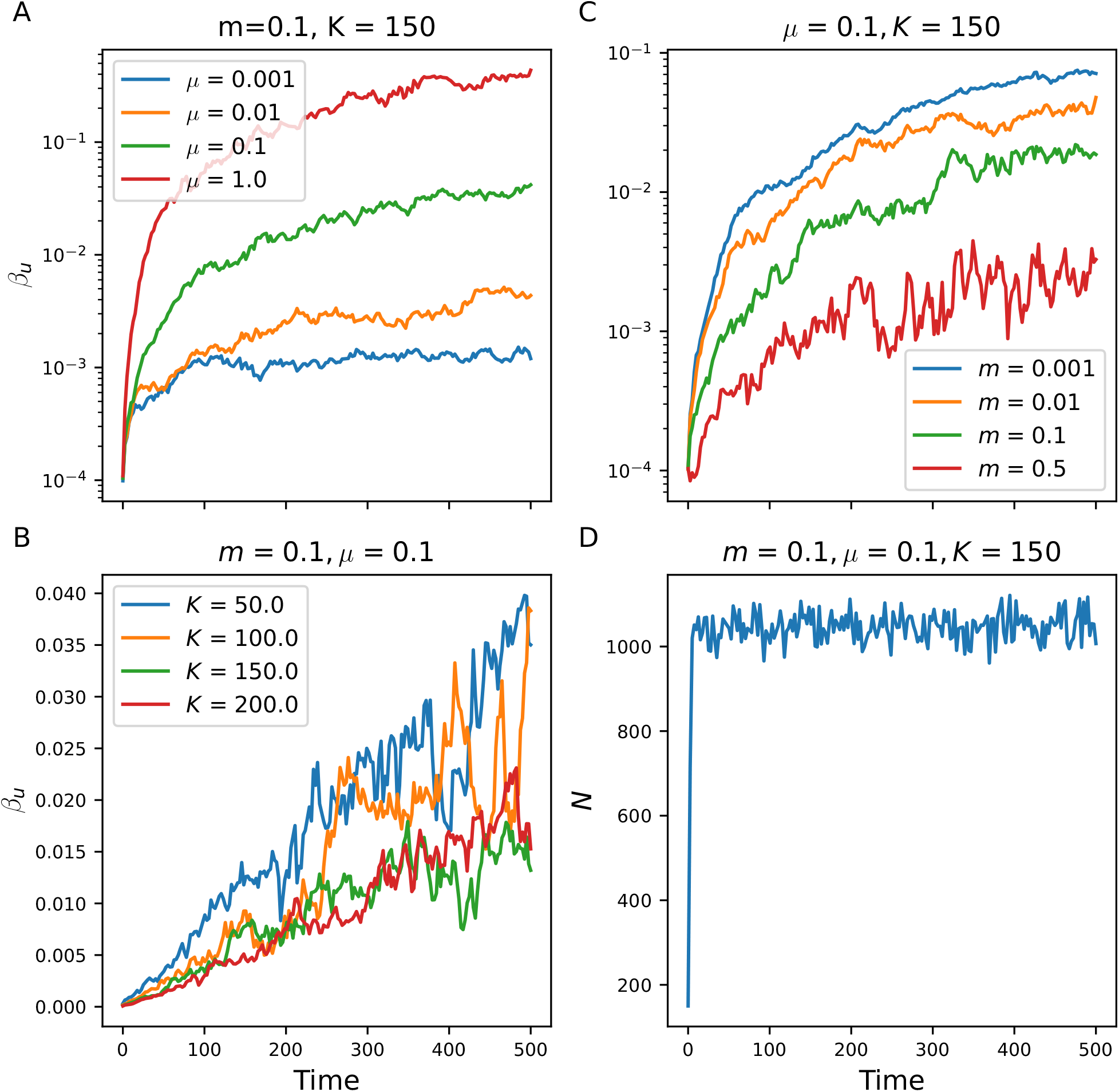
Transient dynamics of *β_u_* and population size in setting (1), for different mutation rates *μ*, migration rates *m*, and carrying capacities *K*. (A) Transient evolution of *β_u_* for varying *μ*. (A) shows that *β_u_* increases with *μ*. (B) Transient evolution of *β_u_* for varying *K*. (B) shows that *β_u_* increases linearly with time at quasi-equilibrium, and illustrates that decreasing K increases *β_u_*, as it favours drift. (C) Transient evolution of *β_u_* for varying *m*. (C) shows that *β_u_* decreases with increasing migration rate. (D) Transient evolution of population size. (D) shows that stochastic fluctuations in population size are negligible.

**Figure S2:**
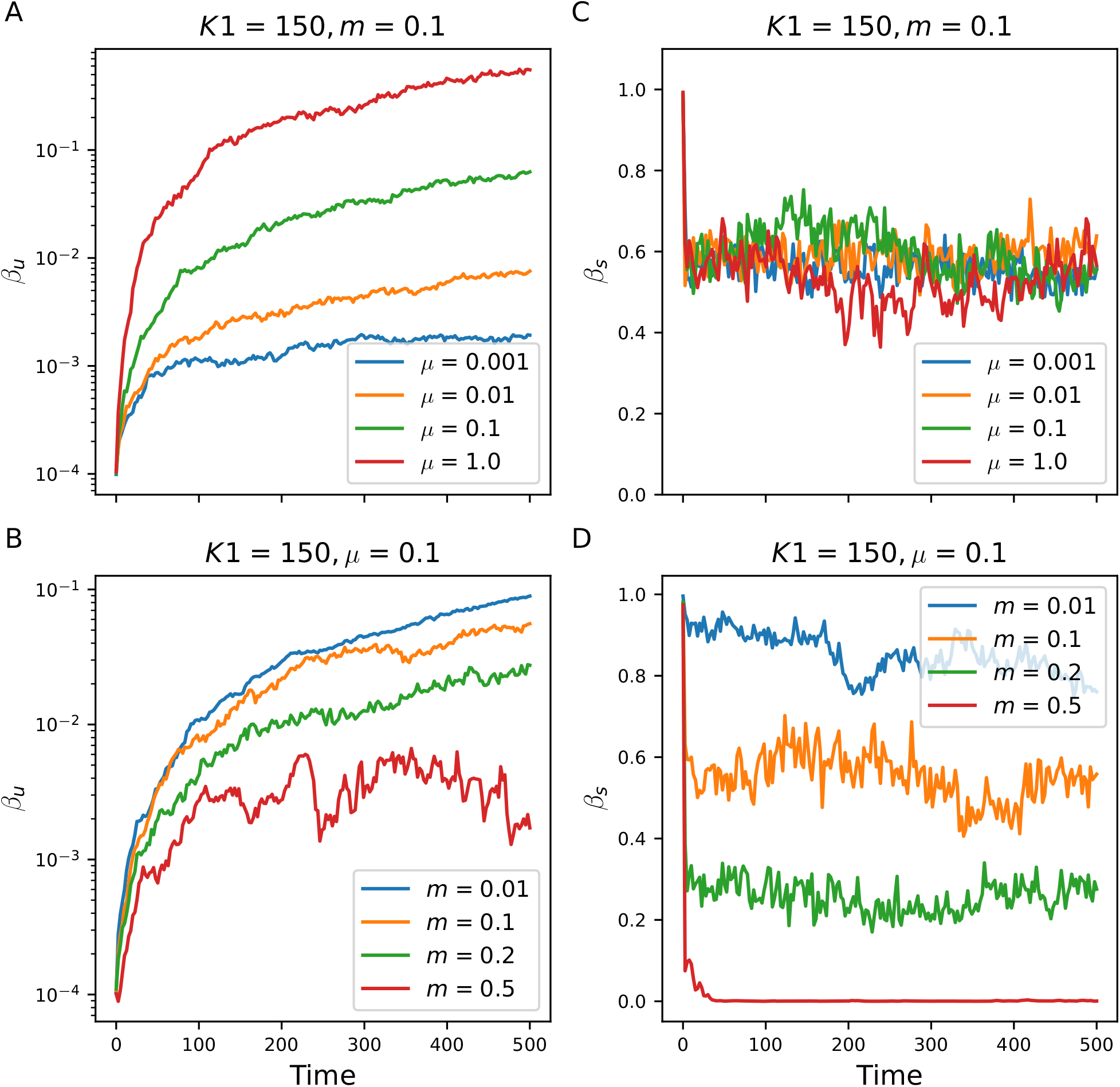
Transient dynamics for *β_u_* and *β_s_* in setting (2), for different mutation rates *μ* and migration rates *m*. (A) and (B) are similar to (A) and (C) in Fig. S1. (C) shows that *β_s_* is not sensitive to *μ*. (D) shows that *β_s_* decreases with increasing *m*. The drop in *β_s_* for *r_θ_* = −1 and *m* = 0.5 corresponds to the case where the population becomes monomorphic.

**Figure S3:**
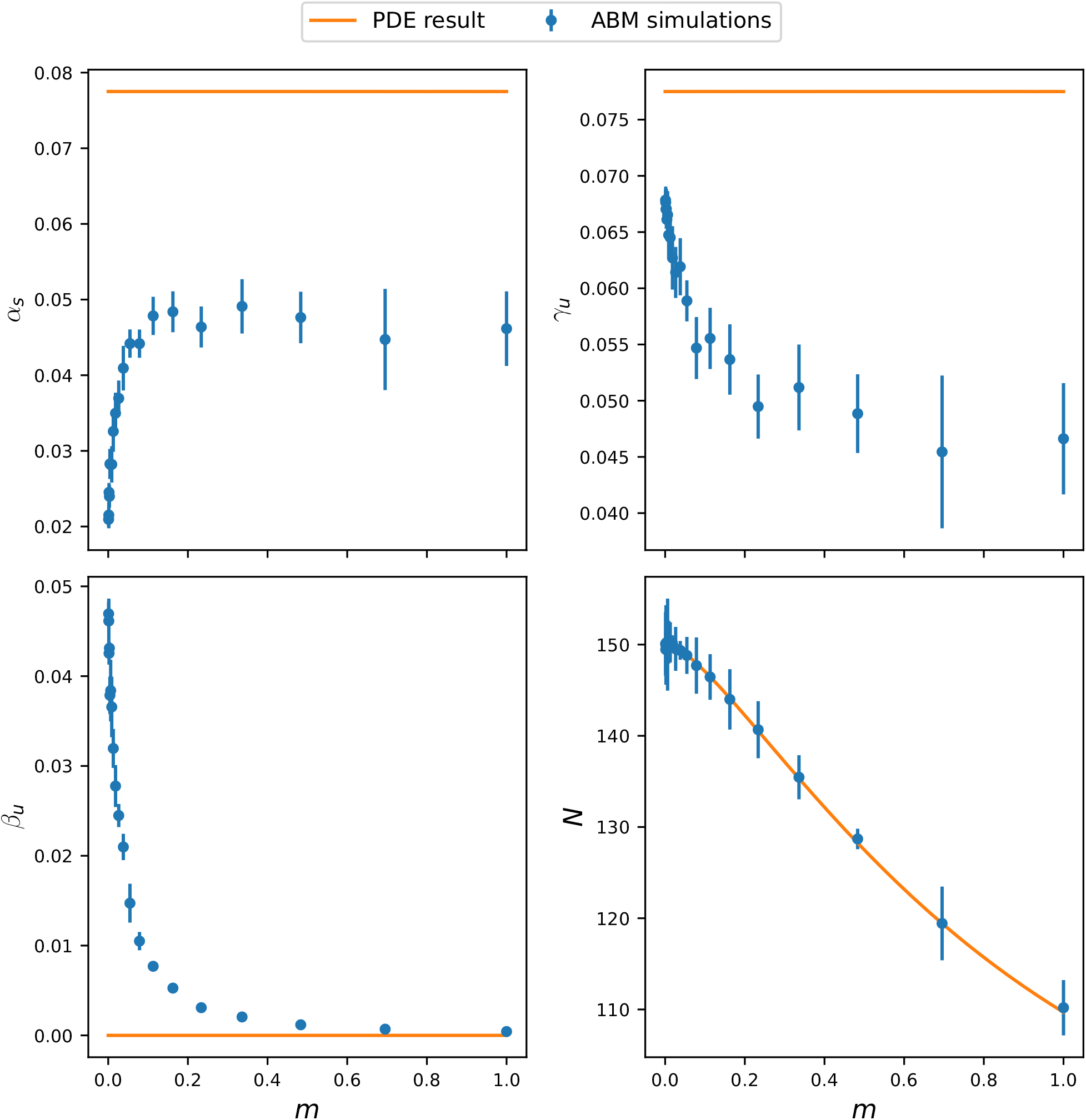
Comparison of neutral diversity metrics and population size obtained from a deterministic approximation (Eqs. (S4) and (S5)) and IBM simulations in setting (1), on a star graph. The figure illustrates that while a deterministic approximation can capture population size, it is not able to capture *α_u_, β_u_* and diversity that characterise different aspect of the neutral trait distribution (see Diversity partitioning for a definition of *α_u_, β_u_* and *γ_u_*).

**Figure S4:**
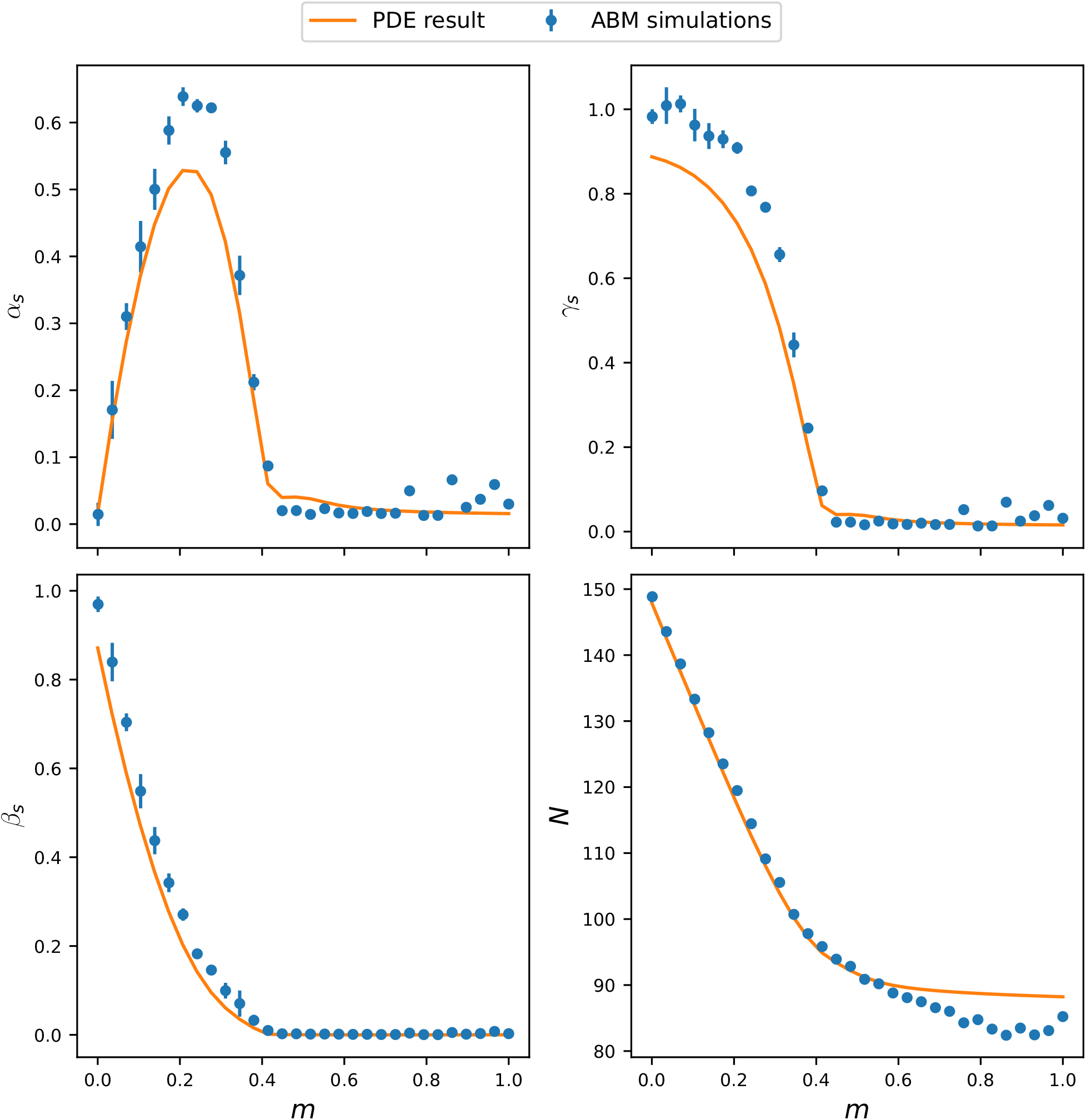
Comparison of adaptive diversity metrics and population size obtained from a deterministic approximation (Eq. (S10)) and IBM simulations in setting (2), on a star graph. The figure illustrates that the metrics *α_s_, β_s_, γ_s_* and population size obtained from the PDE closely match those from the IBM simulations that characterise different aspect of the adaptive trait distribution (see Diversity partitioning for a definition of *α_s_, β_s_* and *γ_s_*).

**Figure S5:**
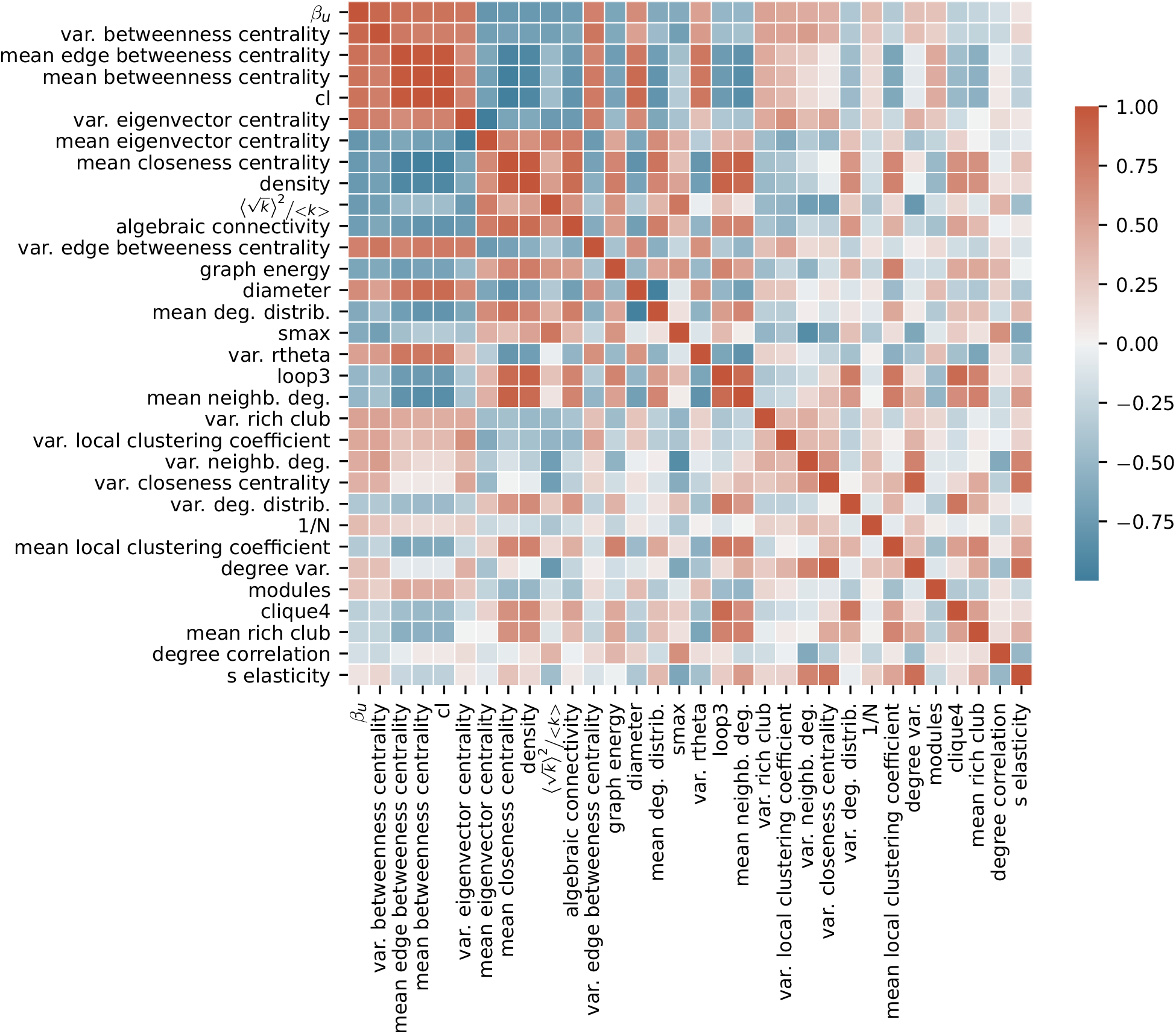
Pearson correlation matrix between *β_u_* and topology metrics for setting (1). On top of a graphical illustration of the correlations provided in Table S1 (column “cor. mean”), the figure shows how topology metrics correlate between each other. Rows are ranked by the absolute value of the correlation between the topological metrics and *β_u_*.

**Figure S6:**
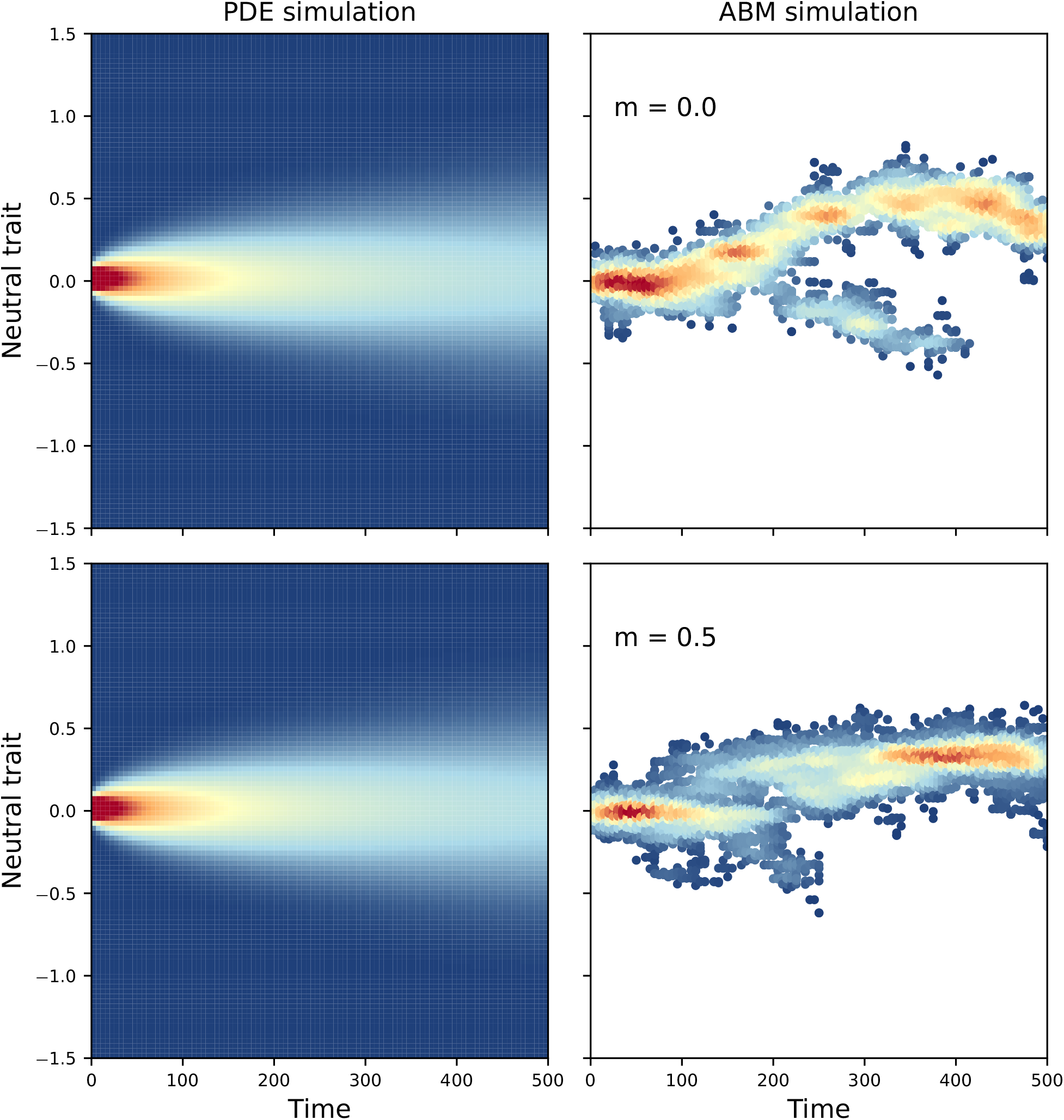
Comparison of the neutral trait density on one vertex obtained from numerically solving Eq. (S5) (left) and IBM simulations (right) in setting (1), for the chain graph. While the PDE model gives a density centred at 0 for *m* = 0, drift arises in the IBM simulation. As *m* grows (lower panels), correlation sets in the local trait pool and the density stabilises to a mean value centred at 0. *K_i_* = 150, *μ* = 0.1, *σ_μ_* = 0.05.

**Figure S7:**
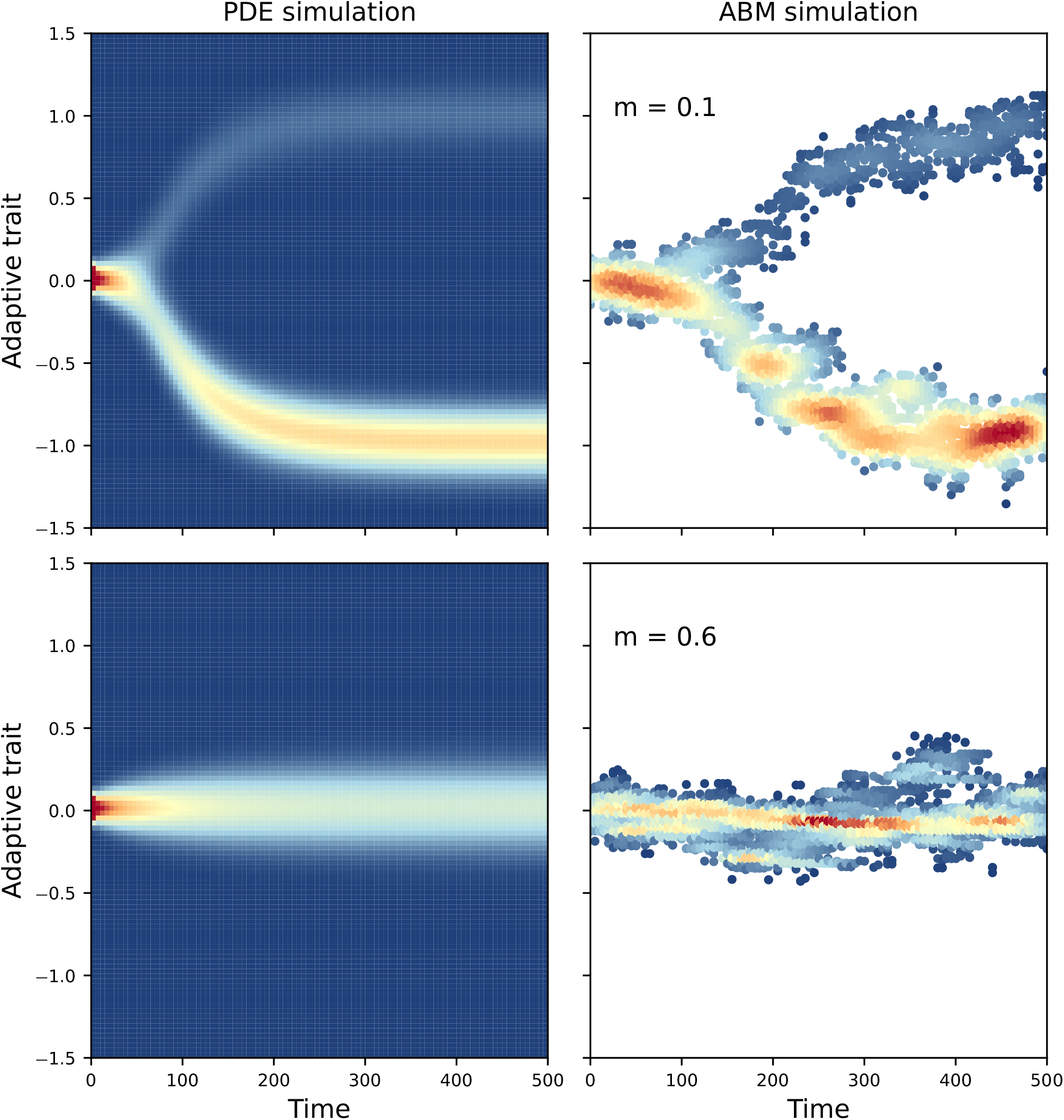
Comparison of the adaptive trait density on one vertex obtained from numerically solving Eq. (S10) (left) and IBM simulations (right) in setting (2), for the star graph. The densities obtained from the IBM and the PDE model look qualitatively similar. *K_i_* = 150, *μ* = 0.1, *σ_μ_* = 0.05, *p* = 0.5.

**Figure S8:**
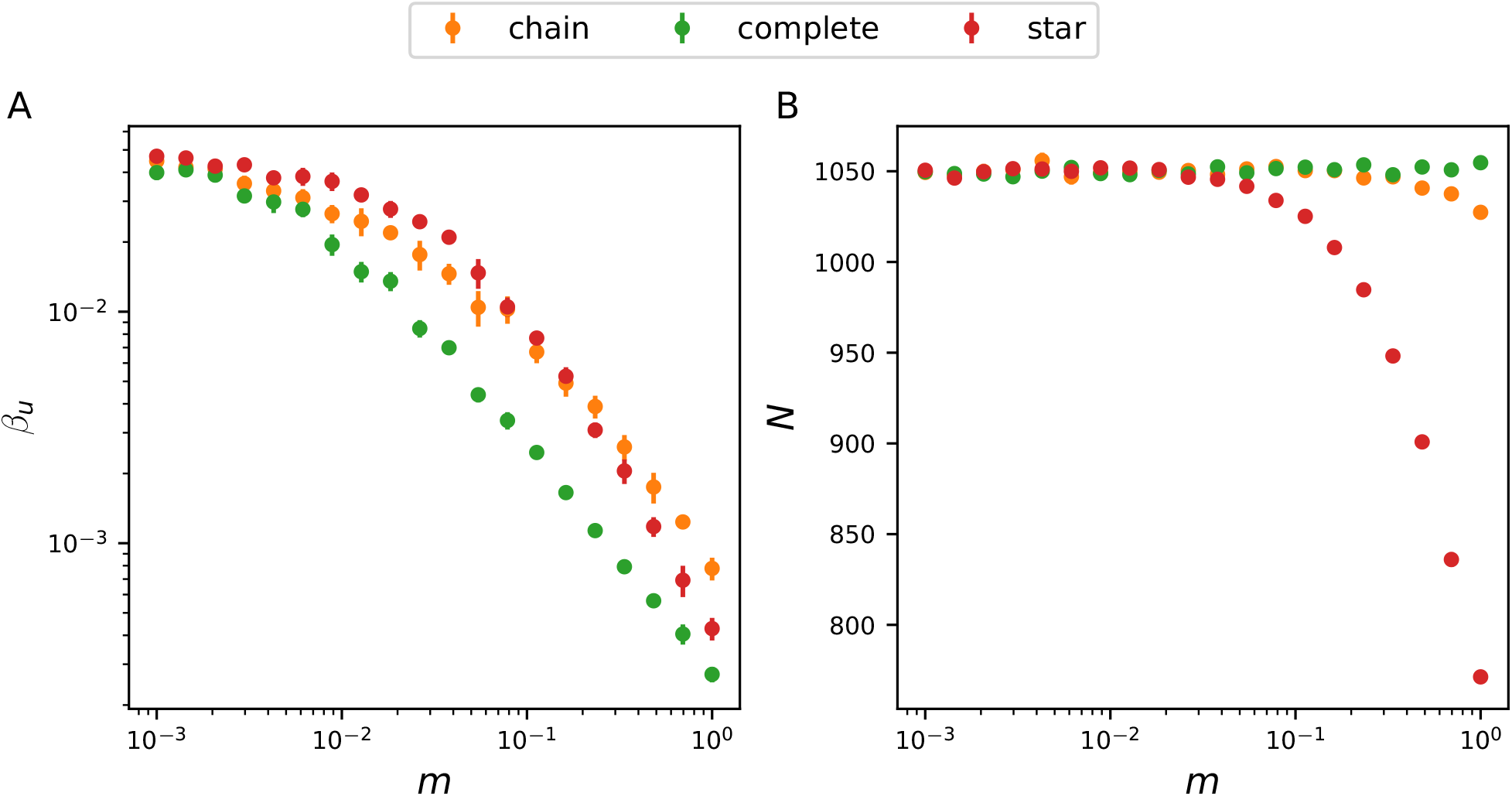
Response of *β_u_* and population size to migration *m* in setting (1), for the chain graph, the complete graph and the star graph. (A) shows that *β_u_* is higher in the star graph for low *m*, but becomes lower than in the chain graph under high migration regimes. (A) therefore suggests a dependency on *m* for the effect of topology metrics on *β_u_*. This dependency appears in Table S1 and Table S3. (B) illustrates that the star graph experiences a large demographic loss under high migration regimes compared with the chain and complete graph.

**Figure S9:**
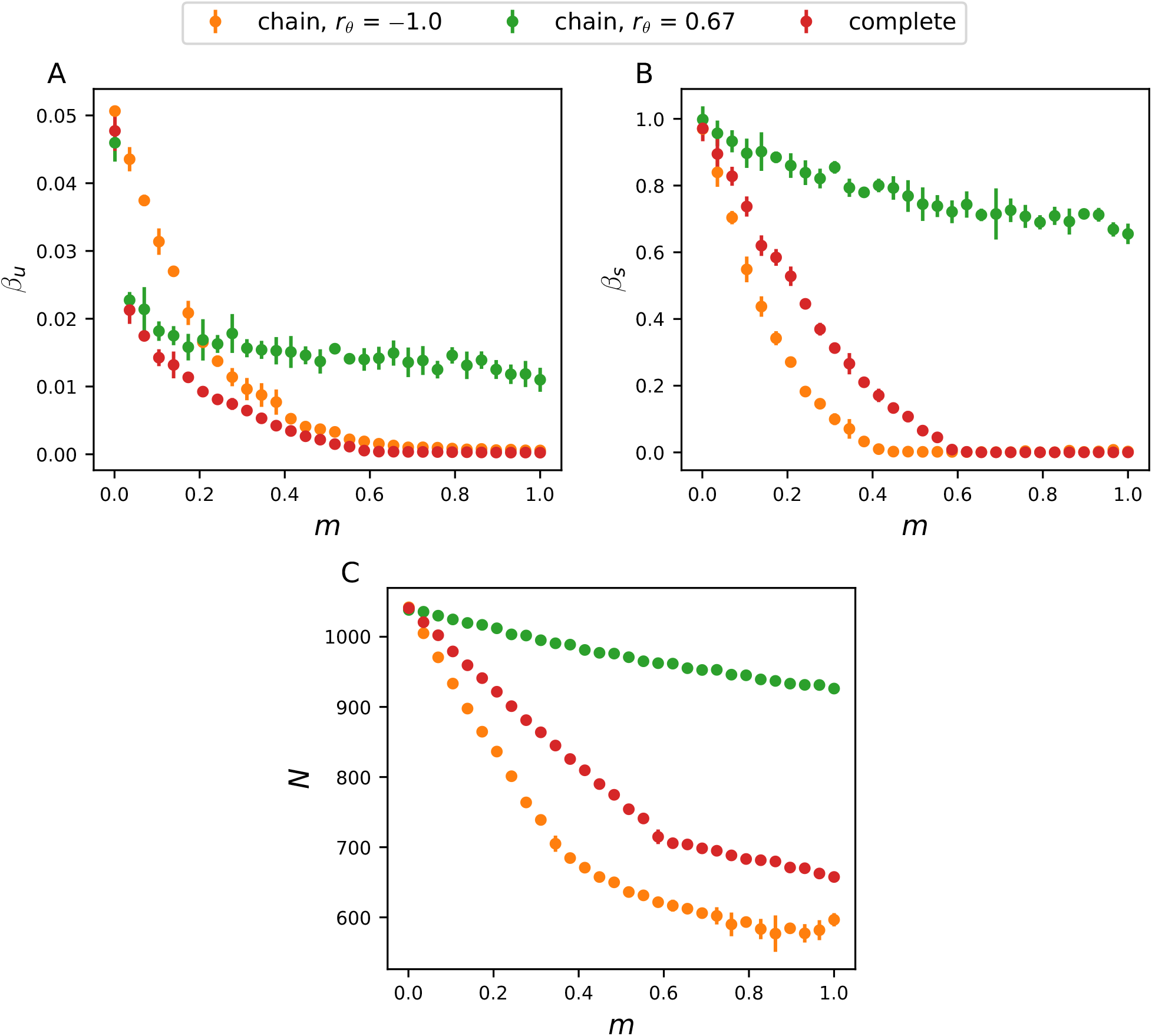
Response of *β_u_, β_s_* and population size to migration *m* in setting (2), for the chain graph with varying *r_θ_* and for the complete graph (*r_θ_* ≈ 0). (A) shows that *β_u_* is higher in the disassortative chain graph for *m* < 0.2, but becomes lower than in the assortative chain graph for *m* > 0.2. (B-C) show that *r_θ_* systematically amplifies *β_s_* and population size irrespective of the migration regime, because populations are better adapted.

**Figure S10:**
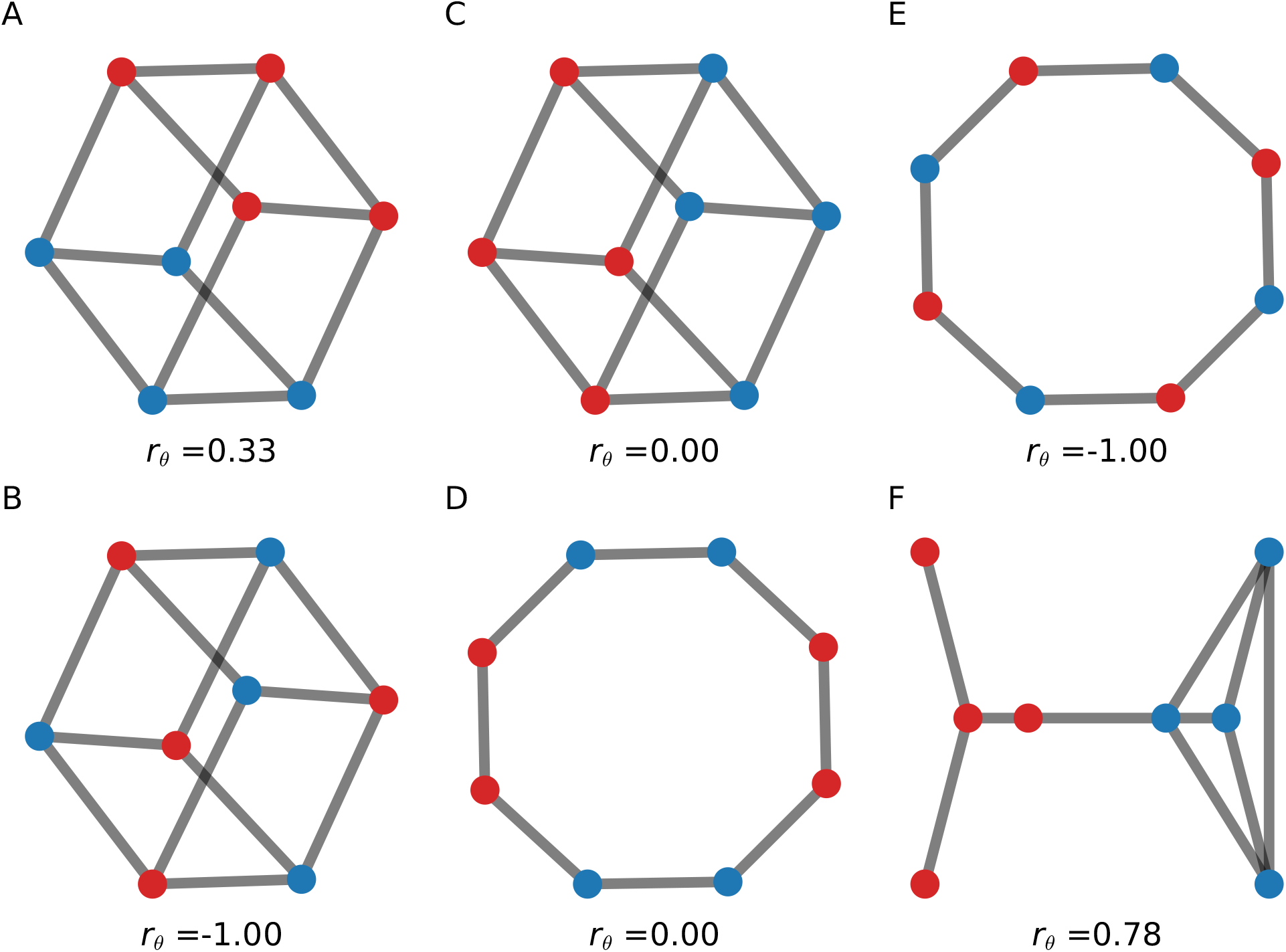
Graphs with different values for habitat assortativity *r_θ_*. Graphs (A-D) can be described exactly with a mean field approach, as blue and red vertices have the equivalent position on the graph.

**Figure S11:**
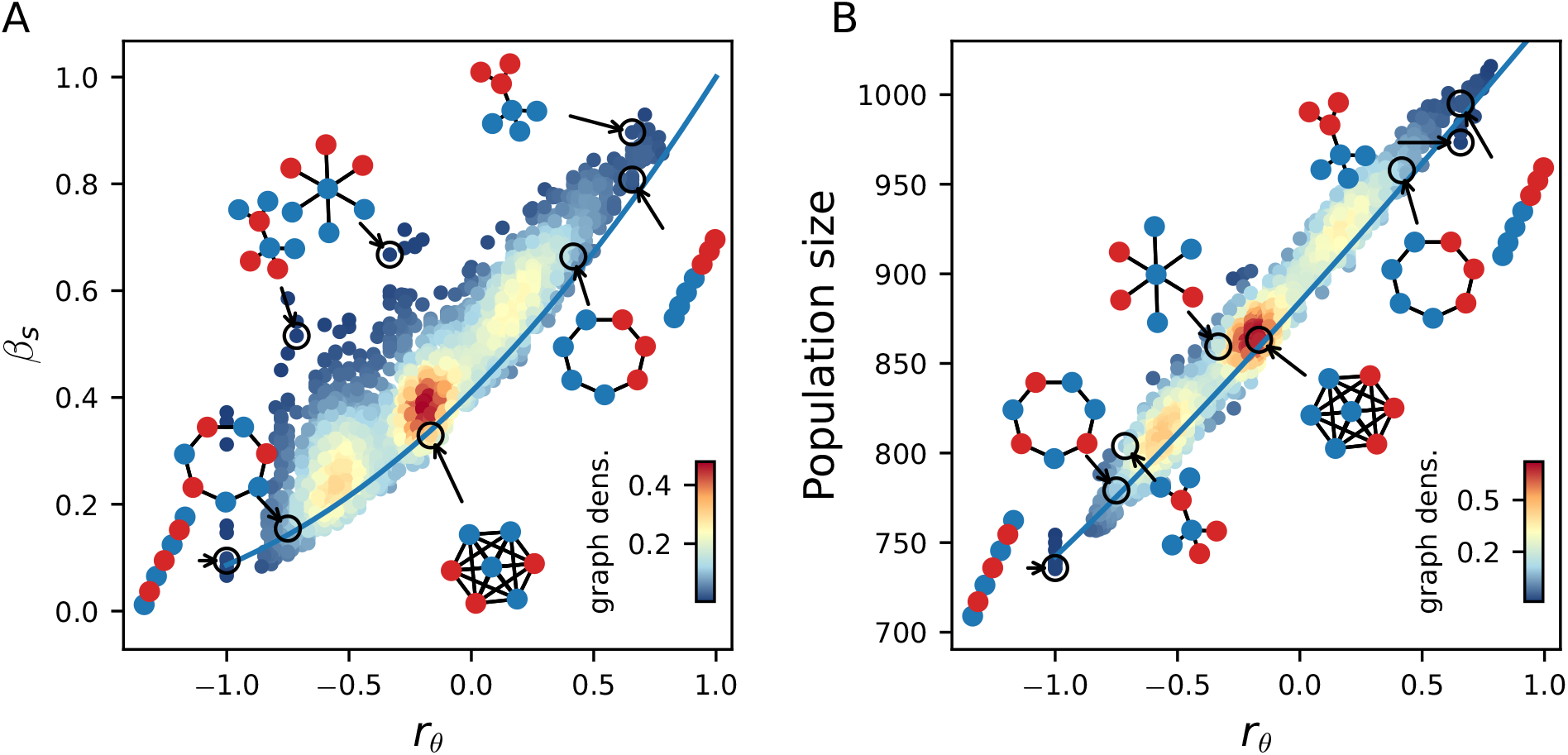
Effect of habitat heterogeneity *r_θ_* on *β_s_* and population size from simulations of the IBM. (A) and (B) show results for simulations on all undirected connected graphs with seven vertices and varying *r_θ_* for *m* = 0.31.

**Figure S12:**
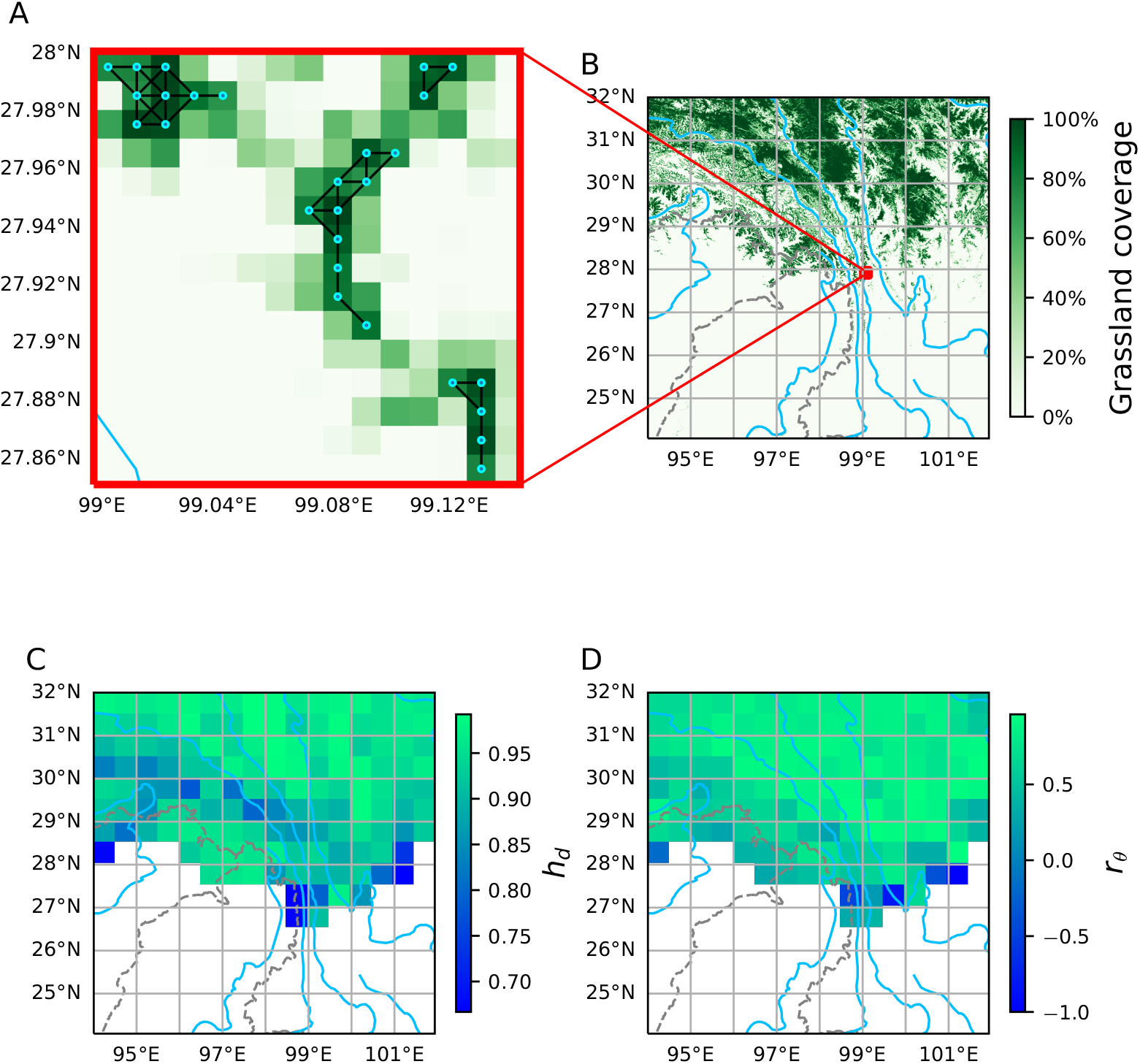
Illustration of the applicability of graph-based metrics to real landscapes. Here we consider a geographical region centered on the Hengduan Moutains in Southwest China, one of the world’s most species-rich temperate alpine biota [4]. (A) Graph representation of a geographical area of size 0.15° × 0.15°. To obtain the graph representation, we consider biological populations living on grasslands, and use the data set provided in [121] containing global grassland coverage at 0.01° resolution. We assign a vertex to a geographical area of size 0.01° × 0.01° if its grassland coverage is above a threshold arbitrarily set to 75%. We further assume that two vertices are connected if the corresponding areas are adjacent, considering that an area is adjacent to its 8 surrounding areas (located at left, top left, top, top right, right, bottom right, bottom, bottom left). (B) Grassland coverage for the considered region. Blue lines correspond to rivers, dashed lines correspond to country borders. (C) and (D) average topology metrics values calculated for the graphs obtained in each geographical area of size 0.25° × 0.25°. (C) Mapping of *h_d_*, the heterogeneity in vertex degree. (D) Mapping of *r_θ_*, the graph assortativity. Annual average temperatures have been considered as the scalar values attributed to each vertex, that capture environmental conditions. Temperature data has been obtained from the Chelsa dataset [122].

**Table S1:** Correlations between *β_u_* and topology metrics for setting (1), ranked by absolute value.

| metrics | $m = 0.01$ | $m = 0.05$ | $m = 0.10$ | $m = 0.50$ | cor. mean | cor. std. |
| --- | --- | --- | --- | --- | --- | --- |
| var. betweenness centrality | 0.83 | 0.90 | 0.90 | 0.88 | 0.88 | 0.03 |
| mean edge betweenness centrality | 0.75 | 0.82 | 0.84 | 0.87 | 0.82 | 0.05 |
| characteristic length | 0.71 | 0.80 | 0.83 | 0.89 | 0.81 | 0.07 |
| var. eigenvector centrality | 0.72 | 0.81 | 0.81 | 0.82 | 0.79 | 0.05 |
| mean eigenvector centrality | -0.71 | -0.81 | -0.81 | -0.82 | -0.79 | 0.05 |
| mean closeness centrality | -0.69 | -0.76 | -0.78 | -0.83 | -0.76 | 0.05 |
| edges density | -0.75 | -0.77 | -0.76 | -0.76 | -0.76 | 0.01 |
| $h_d$ | -0.82 | -0.81 | -0.77 | -0.62 | -0.75 | 0.09 |
| algebraic connectivity | -0.70 | -0.73 | -0.73 | -0.75 | -0.73 | 0.02 |
| var. edge betweenness centrality | 0.57 | 0.71 | 0.76 | 0.87 | 0.73 | 0.12 |
| graph energy | -0.69 | -0.68 | -0.67 | -0.60 | -0.66 | 0.04 |
| diameter | 0.53 | 0.63 | 0.67 | 0.75 | 0.65 | 0.09 |
| mean deg. distrib. | -0.53 | -0.61 | -0.64 | -0.69 | -0.62 | 0.07 |
| $s$ max | -0.73 | -0.67 | -0.61 | -0.45 | -0.61 | 0.12 |
| loop- 3 | -0.53 | -0.51 | -0.50 | -0.51 | -0.51 | 0.01 |
| mean neighb. deg. | -0.42 | -0.48 | -0.52 | -0.61 | -0.51 | 0.08 |
| var. rich club | 0.54 | 0.52 | 0.52 | 0.45 | 0.51 | 0.04 |
| var. local clustering coef. | 0.49 | 0.49 | 0.47 | 0.46 | 0.48 | 0.01 |
| var. neighb deg | 0.59 | 0.49 | 0.43 | 0.29 | 0.45 | 0.13 |
| var. closeness centrality | 0.46 | 0.47 | 0.43 | 0.32 | 0.42 | 0.07 |
| var. deg. distrib. | -0.38 | -0.37 | -0.35 | -0.37 | -0.37 | 0.01 |
| $\frac{1}{N}$ | 0.11 | 0.26 | 0.53 | 0.50 | 0.35 | 0.20 |
| mean local clustering coefficient | -0.35 | -0.34 | -0.36 | -0.34 | -0.35 | 0.01 |
| deg. var. | 0.43 | 0.37 | 0.31 | 0.16 | 0.32 | 0.11 |
| modules | 0.29 | 0.32 | 0.32 | 0.31 | 0.31 | 0.01 |
| clique4 | -0.32 | -0.30 | -0.28 | -0.29 | -0.30 | 0.02 |
| mean rich club | -0.24 | -0.22 | -0.22 | -0.30 | -0.25 | 0.04 |
| deg. correlation | -0.31 | -0.20 | -0.14 | 0.01 | -0.16 | 0.13 |
| $s$ elasticity | 0.21 | 0.13 | 0.07 | -0.06 | 0.09 | 0.11 |

**Table S2:** Correlations between *β_s_* and topology metrics for setting (2), ranked by absolute value.

| metrics | $m = 0.10$ | $m = 0.31$ | cor. mean | cor. std |
| --- | --- | --- | --- | --- |
| $r_\theta$ | 0.94 | 0.91 | 0.93 | 0.02 |
| var. betweenness centrality | 0.19 | 0.33 | 0.26 | 0.10 |
| $h_d$ | 0.17 | 0.29 | 0.23 | 0.09 |
| $s$ max | -0.19 | -0.26 | -0.22 | 0.05 |
| algebraic connectivity | -0.14 | -0.29 | -0.22 | 0.11 |
| var. eigenvector centrality | 0.14 | 0.29 | 0.22 | 0.11 |
| mean eigenvector centrality | -0.14 | -0.29 | -0.21 | 0.11 |
| density | -0.13 | -0.28 | -0.20 | 0.11 |
| mean edge betweenness centrality | 0.12 | 0.28 | 0.20 | 0.11 |
| characteristic length | 0.12 | 0.28 | 0.20 | 0.11 |
| var. edge betweenness centrality | 0.13 | 0.26 | 0.20 | 0.09 |
| mean closeness centrality | -0.11 | -0.27 | -0.19 | 0.11 |
| var. neighb. deg | 0.17 | 0.21 | 0.19 | 0.02 |
| var. local clustering coef | 0.13 | 0.22 | 0.17 | 0.07 |
| var. rich club | 0.13 | 0.20 | 0.17 | 0.05 |
| graph energy | -0.09 | -0.21 | -0.15 | 0.08 |
| var. closeness centrality | 0.12 | 0.17 | 0.14 | 0.04 |
| diameter | 0.07 | 0.21 | 0.14 | 0.10 |
| mean deg. distrib. | -0.07 | -0.21 | -0.14 | 0.10 |
| var. degree | 0.12 | 0.15 | 0.14 | 0.02 |
| loop 3 | -0.07 | -0.18 | -0.13 | 0.08 |
| var. deg. distrib. | -0.08 | -0.16 | -0.12 | 0.05 |
| mean neighb. deg. | -0.04 | -0.17 | -0.10 | 0.09 |
| deg. correlation | -0.10 | -0.09 | -0.09 | 0.01 |
| $s$ elasticity | 0.10 | 0.07 | 0.09 | 0.02 |
| clique 4 | -0.05 | -0.11 | -0.08 | 0.05 |
| mean rich club | -0.03 | -0.08 | -0.05 | 0.03 |
| modules | -0.00 | 0.07 | 0.03 | 0.05 |
| mean local clustering coefficient | 0.01 | -0.07 | -0.03 | 0.06 |

**Table S3:**
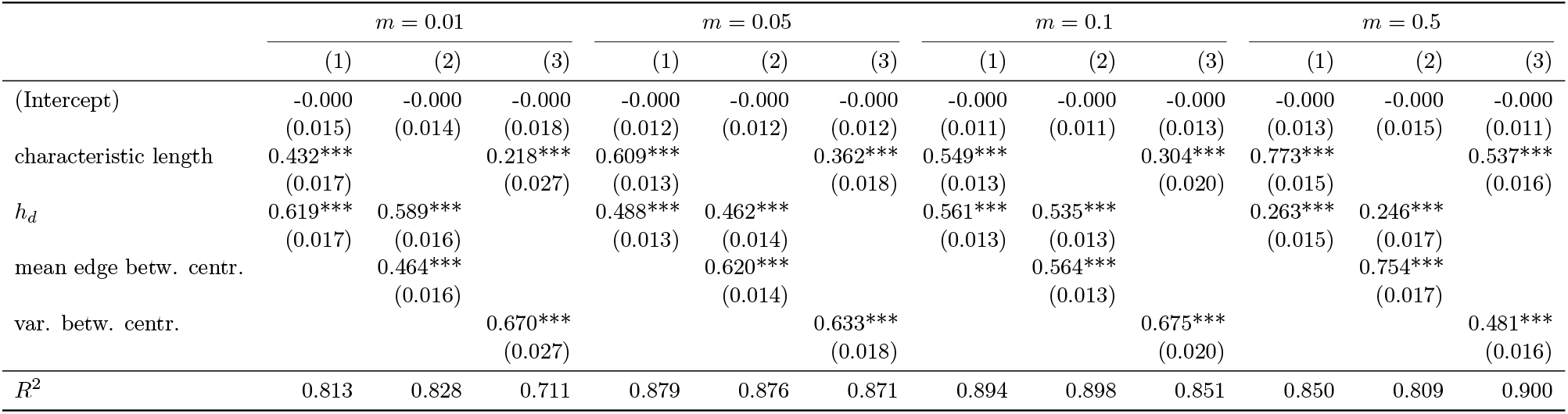
Linear regression model coefficients for the effect of topology metrics on *β_u_*, setting (1). *** *P* < 0.001

| | $m = 0.01$ | | | $m = 0.05$ | | | $m = 0.1$ | | | $m = 0.5$ | | |
| --- | --- | --- | --- | --- | --- | --- | --- | --- | --- | --- | --- | --- |
|  | (1) | (2) | (3) | (1) | (2) | (3) | (1) | (2) | (3) | (1) | (2) | (3) |
| (Intercept) | -0.000<br>(0.015) | -0.000<br>(0.014) | -0.000<br>(0.018) | -0.000<br>(0.012) | -0.000<br>(0.012) | -0.000<br>(0.012) | -0.000<br>(0.011) | -0.000<br>(0.011) | -0.000<br>(0.013) | -0.000<br>(0.013) | -0.000<br>(0.015) | -0.000<br>(0.011) |
| characteristic length | 0.432***<br>(0.017) |  | 0.218***<br>(0.027) | 0.609***<br>(0.013) |  | 0.362***<br>(0.018) | 0.549***<br>(0.013) |  | 0.304***<br>(0.020) | 0.773***<br>(0.015) |  | 0.537***<br>(0.016) |
| $h_d$ | 0.619***<br>(0.017) | 0.589***<br>(0.016) | | 0.488***<br>(0.013) | 0.462***<br>(0.014) | | 0.561***<br>(0.013) | 0.535***<br>(0.013) | | 0.263***<br>(0.015) | 0.246***<br>(0.017) | |
| mean edge betw. centr. |  | 0.464***<br>(0.016) |  |  | 0.620***<br>(0.014) |  |  | 0.564***<br>(0.013) |  |  | 0.754***<br>(0.017) |  |
| var. betw. centr. |  |  | 0.670***<br>(0.027) |  |  | 0.633***<br>(0.018) |  |  | 0.675***<br>(0.020) |  |  | 0.481***<br>(0.016) |
| $R^2$ | 0.813 | 0.828 | 0.711 | 0.879 | 0.876 | 0.871 | 0.894 | 0.898 | 0.851 | 0.850 | 0.809 | 0.900 |

**Table S4:** Linear regression model coefficients for the effect of topology metrics on *β_s_*, setting (2). *** *P* < 0.001

| | $m = 0.1$ | | | $m = 0.31$ | | |
| --- | --- | --- | --- | --- | --- | --- |
|  | (1) | (2) | (3) | (1) | (2) | (3) |
| (Intercept) | -0.000<br>(0.006) | -0.000<br>(0.006) | -0.000<br>(0.006) | 0.000<br>(0.006) | 0.000<br>(0.006) | 0.000<br>(0.007) |
| $h_d$ | 0.170***<br>(0.006) | | | 0.293***<br>(0.006) | | |
| $r_\theta$ | 0.944***<br>(0.006) | 0.935***<br>(0.006) | 0.942***<br>(0.006) | 0.910***<br>(0.006) | 0.893***<br>(0.006) | 0.907***<br>(0.007) |
| var. betw. centrality |  | 0.137***<br>(0.006) |  |  | 0.275***<br>(0.006) |  |
| smax |  |  | -0.180***<br>(0.006) |  |  | -0.247***<br>(0.007) |
| $R^2$ | 0.920 | 0.909 | 0.923 | 0.913 | 0.902 | 0.888 |

**Table S5:**
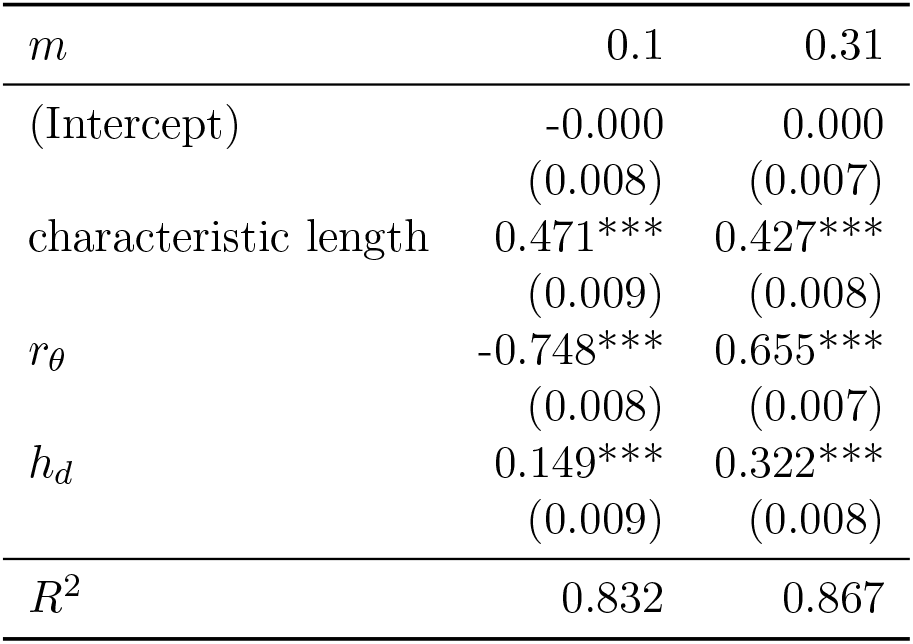
Linear regression model coefficients for the effect of topology metrics on *β_u_*, setting (2). *** *P* < 0.001

## Notes

https://gitlab.ethz.ch/publications/neutral-and-adaptive-diversification-in-spatial-graphs.git

### Competing Interest Statement

The authors have declared no competing interest.

### Summary of Updates

Supplementary Information Text updated to clarify the derivation of the deterministic approximation equations from the baseline stochastic model. Figure S12 added to illustrate the applicability of our theory to real landscapes.

